# Single cell RNA sequencing reveals immunomodulatory effects of stem cell factor and granulocyte colony-stimulating factor treatment in the brains of aged APP/PS1 mice

**DOI:** 10.1101/2024.05.09.593359

**Authors:** Robert S. Gardner, Michele Kyle, Karen Hughes, Li-Ru Zhao

## Abstract

Alzheimer’s disease (AD) leads to progressive neurodegeneration and dementia. AD primarily affects older adults with neuropathological changes including amyloid-beta (Aβ) deposition, neuroinflammation, and neurodegeneration. We have previously demonstrated that systemic treatment with combined stem cell factor (SCF) and granulocyte colony-stimulating factor (G-CSF) (SCF+G-CSF), reduces Aβ load, increases Aβ uptake by activated microglia and macrophages, reduces neuroinflammation, and restores dendrites and synapses in the brains of aged APPswe/PS1dE9 (APP/PS1) mice. However, the mechanisms underlying SCF+G-CSF-enhanced brain repair in aged APP/PS1 mice remain unclear. This study used a transcriptomic approach to identify potential mechanisms by which SCF+G-CSF treatment modulates microglia and peripheral myeloid cells to mitigate AD pathology in the aged brain. After injections of SCF+G-CSF for 5 consecutive days, single-cell RNA sequencing was performed on CD11b^+^ cells isolated from the brains of 28-month-old APP/PS1 mice. The vast majority of cell clusters aligned with transcriptional profiles of microglia in various activation states. However, SCF+G-CSF treatment dramatically increased a cell population showing upregulation of marker genes related to peripheral myeloid cells. Flow cytometry data also revealed an SCF+G-CSF-induced increase of cerebral CD45^high^/CD11b^+^ active phagocytes. SCF+G-CSF treatment robustly increased the transcription of genes implicated in immune cell activation, including gene sets that regulate inflammatory processes and cell migration. Expression of S100a8 and S100a9 were robustly enhanced following SCF+G-CSF treatment in all CD11b^+^ cell clusters. Moreover, the topmost genes differentially expressed with SCF+G-CSF treatment were largely upregulated in S100a8/9-positive cells, suggesting a well-conserved transcriptional profile related to SCF+G-CSF treatment in resident and peripherally derived CD11b^+^ immune cells. This S100a8/9-associated transcriptional profile contained notable genes related to pro-inflammatory and anti-inflammatory responses, neuroprotection, and Aβ plaque inhibition or clearance. Altogether, this study reveals immunomodulatory effects of SCF+G-CSF treatment in the aged brain with AD pathology, which will guide future studies to further uncover the therapeutic mechanisms.

## Background

Alzheimer’s disease (AD) is the most prominent neurodegenerative disease and the leading cause of dementia [1, 2]. The primary risk factor for AD is advanced age. Strikingly, approximately one in nine Americans over the age of 65 is currently living with AD-related dementia. Moreover, societal impacts of AD are immense with annual healthcare costs greater than $320 billion [2]. Without novel effective treatments, the number of Americans afflicted with AD is projected to reach 13.8 million by 2060 [3].

Neuropathological hallmarks of AD include extracellular plaques composed of aggregated amyloid-beta (Aβ), intracellular neurofibrillary tangles composed of aggregated tau, and neuroinflammation [4, 5]. Aβ peptides generated by sequential proteolysis of amyloid precursor protein (APP) by β- and γ-secretases are particularly prone to aggregation [6, 7], resulting in plaques that contribute to tau pathology, neuro-inflammation, synaptic dysfunction, neurodegeneration, and cognitive decline [8, 9, 10, 11, 12, 13].

Several studies crossing Aβ and tau transgenic animal models that develop extracellular plaques and intracellular tangles, respectively, revealed that while Aβ pathology enhanced the formation of neurofibrillary tangles, tau pathology had relatively little effect on Aβ deposits [14, 15, 16]. Injections of Aβ fragments, oligomers, or fibrils into the brain are shown to enhance the formation of neurofibrillary tangles [17], increase neuroinflammation and neurodegeneration, and impair spatial learning and memory [18, 19, 20]. These collective findings suggest that Aβ neuropathology is a contributory factor and relatively early event in the progression of AD, and that Aβ clearance is a promising focus for therapeutic advancements [21]. Indeed, clearance of Aβ from the brain is associated with mitigation of neurodegeneration and cognition decline, largely observed in mouse models [22, 23, 24].

Effective and safe treatments that can stop or delay pathological progression for AD patients, however, are not currently available. Approved and prescribed treatments for AD were primarily symptomatic, for example, targeting behavioral problems, rather than the underlying neuropathology, and thus, do not stop or reverse the progression of the disease [25]. Anti-Aβ monoclonal antibodies have been approved by the United States Food and Drug Administration for treatment of early AD patients presenting with Aβ pathology and have been shown to reduce Aβ and improve cognitive function; however, concerns over Aβ antibody treatments include high treatment costs, accessibility, and the incidence of adverse events including brain edema, microhemorrhages, and brain volume loss [26–28]. It remains highly imperative to develop safe and effective treatments for AD patients.

Microglia are the primary phagocytic cells in the brain. They act as the first line of immune defense, surveilling the microenvironment, clearing debris or pathogens, including aggregated Aβ plaques [29], to maintain homeostasis [30]. In AD, bone marrow-derived blood myeloid cells, most notably monocytes, augment the brain’s immune cell population by migrating to the brain and differentiating into macrophages, also showing robust efficacy to uptake and degrade aggregated Aβ [31, 32, 33]. Notably, mutations in genes related to microglial and macrophage activation, such as triggering receptor expressed on myeloid cells 2 (Trem2), a transmembrane receptor activated by Aβ [34] and associated with Aβ clearance [35, 36], confer relatively high risk of AD [37, 38].

Microglia and macrophages that migrate to and surround Aβ plaques release a host of extra-cellular vesicles and soluble molecules, including cytokines with inflammatory and/or anti-inflammatory functions, and thus, additionally regulate neuroinflammatory processes [39, 40]. Activated microglia and macrophages associated with a relatively anti-inflammatory profile are thought to confer debris clearance without significantly contributing to inflammation and may enhance angiogenesis and tissue repair [41, 42, 43]. However, during the progression of AD pathology, microglia and macrophages increasingly take on a relatively pro-inflammatory state [44, 45, 46]. Although numerous reports demonstrate Aβ clearance is enhanced by pro-inflammatory signaling or cytokines [23, 47, 48, 49, 50], persistent or chronic inflammation is generally thought to enhance Aβ spreading and aggregation [51], creating a vicious cycle leading to neurodegeneration [52]. Altogether, these previous studies strongly suggest that the development of novel AD therapies to clear Aβ from the brain, should consider their effects on inflammation.

Stem cell factor (SCF) and granulocyte colony-stimulating factor (G-CSF) are two hematopoietic growth factors that synergistically stimulate proliferation, differentiation, and mobilization of hematopoietic stem cells and progenitor cells, significantly increase the population of blood leukocytes, and augment the immune response [53, 54, 55, 56]. SCF and G-CSF also act directly on neurons, glia, and blood vessel cells [57, 58, 59], in the central nervous system. SCF and G-CSF treatment enhances angiogenesis, neural survival, neurite outgrowth, synaptogenesis, and neurogenesis, and reduces neuroinflammation in several models of neurodegenerative diseases, neurological disorders, and brain trauma [60, 61, 62, 63, 64, 65, 66]. In AD patients, plasma levels of SCF and G-CSF are significantly decreased [67, 68]. Clinical studies have also demonstrated inverse correlations between the plasma levels of SCF or G-CSF and AD severity [68, 69] and Aβ levels in cerebrospinal fluid [70].

Demonstrating the efficacy of SCF and G-CSF to treat AD neuropathology in pre-clinical studies, our lab has observed that systemic treatment of combined SCF and G-CSF (SCF+G-CSF) leads to long-lasting reductions of Aβ plaques in the hippocampus and cortex of middle-aged [71] and aged [22] APP/PS1 mice, a commonly used mouse model of Aβ pathology in AD research. Coincident with these findings, SCF+G-CSF treatment increases the association between Aβ plaques and activated Trem2^+^ microglia and macrophages, increases Aβ contained in CD68^+^ lysosomal compartments, increases the density of the anti-inflammatory marker IL-4, and decreases the density of the pro-inflammatory marker NOS-2 in the hippocampus and cortex of aged APP/PS1 mice [22]. These collective findings suggest that SCF+G-CSF treatment enhances Aβ clearance by activating microglia and macrophages to uptake and degrade Aβ plaques, while also shifting the environment toward a relatively anti-inflammatory state. Importantly, these effects further correspond to SCF+G-CSF-mediated reductions of aggregated tau and increases in dendritic marker MAP2 and post-synaptic marker PSD-95, suggesting treatment-related rebuilding of neural connections [22]. Thus, SCF+G-CSF treatment in aged APP/PS1 mice ameliorates or reverses each central feature of AD neuropathology: aggregated Aβ, aggregated tau, neuroinflammation, and degeneration of neural processes and synaptic connections [22]. These findings indicate that SCF+G-CSF treatment changes the functions of microglia and macrophages to mitigate AD neuropathology in the aged brain.

Leveraging a single-cell RNA sequencing approach, the aim of the present study is to identify novel transcriptional profiles of microglia and myeloid cells in the brains of aged APP/PS1 mice following SCF+G-CSF treatment. Similar approaches have been used to identify novel functional states of microglia related to AD, including a profile of disease associated microglia (DAM) found to limit the progression of AD pathology [72, 73]. Here, we analyzed the transcriptional profiles of CD11b^+^ microglia and myeloid cells isolated from the brains of aged APP/PS1 mice treated with SCF+G-CSF or vehicle solutions. We profiled transcriptional changes of brain CD11b^+^ cells the day after a 5-day treatment to identify SCF+G-CSF-induced transcriptional responses. The findings of this study provide unbiased identification of SCF+G-CSF treatment-related transcriptional profiles of immune cells in the brain, which may guide future mechanistic studies to understand how SCF+G-CSF treatment mitigates AD pathology.

## Materials and Methods

### Animals

All methods were carried out in accordance with the National Institutes of Health Guide for the Care and Use of Laboratory Animals and approved by the State University of New York Upstate Medical University Institutional Animal Care and Use Committee. Aged APP/PS1 mice (male, 28 months old) (stock# 034832, Jackson Labs) were used in these experiments. APP/PS1 mice express both chimeric amyloid precursor protein (human APP695swe) with Swedish double mutations (K595N/M596L) and human presenilin protein 1 carrying the exon-9-deleted variant (PS1-dE9) [74]. APP/PS1 mice develop plaques of human amyloid beta (Aβ) peptide in the brain by 6-7 months of age. Mice had free access to food and water and were housed in a 12-hour light/dark cycle. The health status of the mice was checked daily.

### Experimental Design

The experimental design is summarized in Figure 1. Twelve APP/PS1 mice at the age of 28-months old were subcutaneously injected with stem cell factor (SCF; 200 µg/kg in saline) and granulocyte colony-stimulating factor (G-CSF; 50 µg/kg in 5% dextrose) or an equal volume of vehicle solution (n=6 in each group) for five consecutive days. On the morning of day 6 after completing the 5-day injections, whole brains (3 mice in each group) were removed and dissociated. CD11b^+^ cells were isolated using magnetic-activated cell sorting (MACS). The isolated CD11b^+^ cells from the brains of 3 mice in the same treatment group were pooled together for single-cell RNA sequencing (scRNAseq). The isolated CD11b^+^ cells were also analyzed using flow cytometry for immuno-phenotyping. Flow cytometry was performed on aliquots of the pooled CD11b^+^ cells used for scRNAseq and on CD11b^+^ cells of individual samples isolated from the brains of 6 additional mice (3 in each group).

**Fig. 1.**
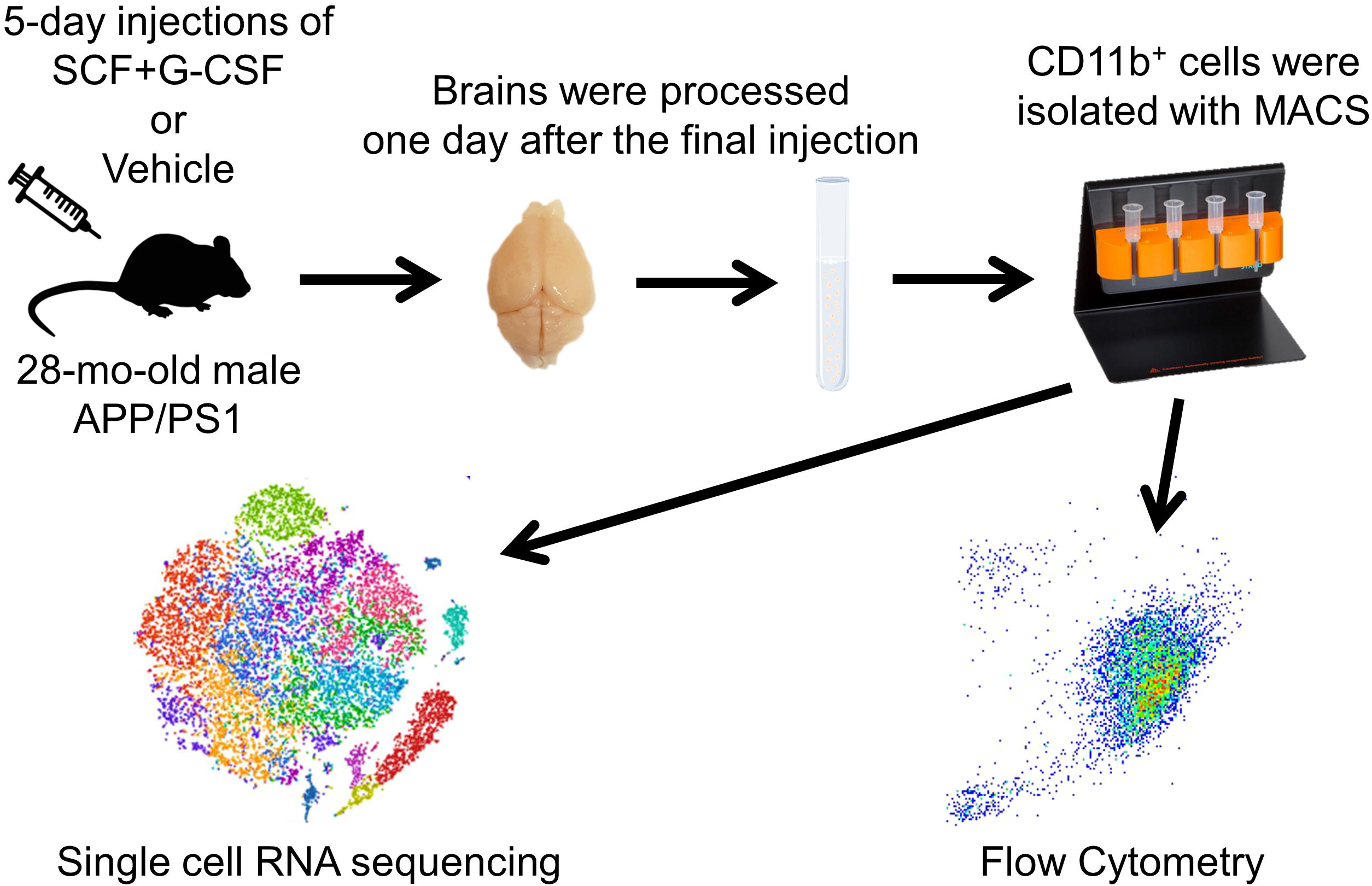
Experimental flow chart. APP/PS1 mice (28-month-old; male) were subcutaneously injected with either combined stem cell factor (SCF) and granulocyte colony-stimulating factor (G-CSF) or vehicle solution for 5 days. The next day, the whole brains were excised and processed into single-cell suspensions. After debris and dead cells were removed, CD11b^+^ cells were isolated using magnetic-activated cell sorting (MACS). The isolated CD11b^+^ cells were used for either single-cell RNA sequencing or flow cytometry.

### Cell processing and CD11b^+^ isolation for single cell RNA sequencing and flow cytometry

#### Cell dissociation

Mice were anesthetized with Ketamine (100mg/kg) and Xylazine (10mg/kg) (i.p.) and euthanized by intracardiac perfusion with ice-cold heparinized (10k U / L; Sagent Pharm. 25021-403-04) Dulbecco’s phosphate-buffered saline (D-PBS; 14190136; ThermoFisher). Each brain was removed, immersed in a cell culture dish containing ice-cold D-PBS+ (D-PBS supplemented with calcium, magnesium, glucose and pyruvate; 14287080; ThermoFisher), and minced (∼1-2 mm in any dimension) using an ice-cold sterile scalpel blade. Brains were dissociated using the commercially-available Adult Brain Dissociation Kit (130-107-677; Miltenyi Biotec) and the Gentle MACS Octo Dissociator with Heaters (130-096-427; Miltenyi Biotec) according to manufacturer specifications. Briefly, tissue pieces from each brain were transferred into a C-tube (130-096-334; Miltenyi Biotec) containing 1950 µL of enzyme mix 1. Enzyme mix 2 was subsequently added (30 µL), and the sample was dissociated using the program 37C_ABDK_01. Each sample was spun down, and the re-suspended pellet was passed through pre-wet 400 µM (43-50400-03; pluriSelect) and 70 µM cell filters (43-50070-51; pluriSelect). Each cell suspension was centrifuged at 300 x g for 10 min at 4°C, and the cell pellet was re-suspended in 3.1 mL D-PBS+ for subsequent processing to clear debris and dead cells.

#### Debris and dead cell removal

Debris removal solution (900 µL; 130-109-398; Miltenyi Biotec) was mixed into each suspension and 4 mL of D-PBS+ was layered over top. The samples were centrifuged at 3000 x g for 10 min at 4°C.The top and debris inter-phases were removed, and the remaining solution was diluted in D-PBS+ (14 mL), inverted three times, and centrifuged at 1000 x g for 10 min at 4°C. Dead and dying cells were removed using the Dead Cell Removal kit from Miltenyi Biotec (130-090-101). Briefly, the pellets were resuspended and incubated for 15 min at room temperature in 100 µL of the dead cell removal magnetically-labeled antibody. Binding Buffer (400 µL) was added to the suspension, which was subsequently passed through a pre-washed LS column (130-042-401; MB; 1 LS column per brain), placed in a strong magnetic field (Quadro MACS Separator,130-090-976, attached to the MACS MultiStand, 130-042-303; Miltenyi Biotec), and topped with a pre-wet 70 µM cell filter. Each filter/column was further washed 4 times with Binding Buffer. Flow-through, largely depleted of dead cells (confirmed by trypan blue staining of cell aliquots), was collected and centrifuged at 300 x g for 10 min at 4°C. Each pellet was resuspended in 1 mL 1X MACS buffer (130-091-376; Miltenyi Biotec; diluted in PBS) and passed through a 40 µM cell filter (43-50040-51; pluriSelect). Additional MACS buffer (4 mL) was passed through the 40 µM filter to maximize cell recovery. The suspension from each brain was centrifuged at 300 x g for 10 min at 4°C, and each pellet was resuspended in 270 µL MACS buffer prior to magnetic activated cell sorting of CD11b^+^ brain cells.

#### Magnetic-activated cell sorting of CD11b^+^ brain cells

CD11b ultra-pure micro beads (30 µL; 130-126-725; Miltenyi Biotec) were incubated with the sample for 15 min at 4°C. Subsequently, each cell suspension was washed with 2 mL MACS buffer and centrifuged at 300 x g for 10 min at 4°C. Each pellet was resuspended in 500 µL MACS buffer and passed through a pre-washed LS column (1 LS column per brain) placed in the magnetic field. Each LS column was topped with a pre-wet 70 µM cell filter. Each filter/column was further washed 3 times with MACS buffer. To enhance CD11b purity, the magnetically-captured cells were eluted into a second LS column affixed to the magnet. The second LS column was washed 3 times with MACS buffer. Viability of magnetically-captured cells (eluted into a new tube away from the magnet) was assessed by trypan blue staining (1:2 dilution sample in trypan blue; BioRad 1450013). Cell viability was quantified as the proportion of cells that did not take up the dye. Aliquots were further processed for scRNAseq and for flow cytometry.

### Single-cell RNA sequencing

The cell suspensions were centrifuged at 300 x g for 10 min at 4°C, and the pellets were resuspended in DMEM/F12 (11320033; ThermoFisher) supplemented with Fetal Bovine Serum (10%) to achieve a concentration of 1000 cells / µL. The cell solutions were placed on ice and run for scRNAseq using the Chromium single cell gene expression platform (10x Genomics) to target 10,000 cells. Following manufacturer specifications (GEM Single Cell 3’ Reagent Kit v3.1: 10x Genomics), samples and barcoded gel beads were loaded onto a G chip and run on a Chromium Controller to partition single cells for generation of cell-specific barcoded cDNA. Pooled cDNA was sequenced using the Illumina NextSeq 500 High Output Kit. The single-cell RNA sequencing was performed by the Molecular Analysis Core Facility at SUNY Upstate Medical University.

### Flow cytometry

Separate aliquots of cell suspensions used for scRNAseq were centrifuged at 300 x g for 10 min at 4°C, and the pellets were resuspended in D-PBS supplemented with 1% BSA. Aliquots (100 µL) were incubated with anti-mouse CD11b-APC (1:50 dilution; 130-113-802 Miltenyi Biotec) and anti-mouse CD45-PE (1:50 dilution; 130-110-797, Miltenyi Biotec) for 10 min at 4°C. Separate aliquots were incubated with control isotype antibodies. Cells were washed with 2 mL D-PBS (1% BSA) and centrifuged at 300 x g for 10 min at 4°C. The pellets were resuspended in 600 µL D-PBS (1% BSA) for flow cytometry analysis. Additional samples were stained and analyzed as above using the following modifications: the samples were blocked with FcR block (1:200 dilution; 553141, BD Biosciences) for 10 min at 4°C and incubated with anti-mouse CD11b-PE (1:50 dilution; 130-113-806, Miltenyi Biotec) and anti-mouse CD45-APC (1.15:100 dilution; 103112, Biolegend) for 15 and 30 min, respectively at 4°C. SCF+G-CSF-treated and vehicle-control samples were counterbalanced across all experiments. Cells were assayed on the BD LSR-II cytometer with FACSDiva software (BD Biosciences) using 561nM and 633nM excitation lasers paired with detection filters 585/15nM and 630/20nM for PE and APC signals, respectively.

### Data analysis and statistics

#### Single cell RNA sequencing

Analysis and figure generation of scRNAseq data were primarily performed using Partek Flow software. Reads were trimmed and aligned to the mm10 assembly using Bowtie2. To ensure the results presented here were robust to choice of aligners, primary findings were corroborated using alternate aligners, including STAR. To limit artifacts, unique molecular identifiers (UMIs) that identify individual transcripts were de-duplicated. Moreover, cell-specific barcodes were filtered to limit those not associated with a cell, using default parameters. A single cell count matrix was generated by quantifying the reads for each cell barcode to mm10_refseq_v97 annotations. Based on recommendations for using 10x genomics kits, the minimum read overlap for inclusion in the count matrix was set to 50% of read length. Strand specificity was set to Forward-Reverse. Low quality cells (e.g., doublets, those with few reads) were manually filtered out based on the distributions of read counts, detected genes, and ribosomal counts across the entire dataset. Manual filtering was done blindly with respect to treatment condition and individual gene profiles. This method yielded a total of 19,008 cells. Normalization was performed to account for differences in total UMI counts per cell and values were log2-transformed. To reduce noise and limit analysis of genes with little-to-no expression, genes with counts less than 1 in 99% of cells were excluded from analysis. Data were dimensionally reduced by Principal Component Analysis. The ten principal components explaining the most variance in the dataset were used to identify clusters. Unsupervised cluster analysis was performed using the Louvain algorithm. t-Distributed stochastic neighbor embedding (tSNE) dimensional reduction plots were generated to visualize cell clusters and their associated gene expression profiles.

To evaluate changes across treatment groups (SCF+G-CSF vs. Vehicle) in cell distributions across clusters, chi-square analysis was performed (Graph Pad v9) with alpha set to 0.05. Differentially expressed genes across conditions were identified using the Biomarker function in Partek. For restricted gene sets (e.g., homeostatic or DAM gene sets), differentially expressed genes showing a ≥ 0.4 log2-fold change across conditions, with a false discovery rate (FDR)-adjusted (step-up method) p value < 0.05 [75] were highlighted as significant. For analyses across the entire transcriptome, significance thresholds were set to a ≥ log2-fold change across conditions with an FDR-adjusted p value < 0.01. This threshold limited significant hits to those genes most robustly altered by SCF+G-CSF treatment. The bioinformatics tool VolcaNoseR [76] was used to generate volcano plots of differentially expressed genes. Enriched gene ontologies, Reactome pathways, and gene interaction networks (generated by querying the STRING database) were identified and analyzed using default settings in Cytoscape 3.9.1 [77, 78].

#### Flow Cytometry

Data processing was performed using Flow Jo v10. Debris was gated out using forward and side scatter profiles, and single cells were selected based on forward scatter area by forward scatter height (Fig. 2A-B). Isotype-stained controls were used for thresholding and identification of CD11b^+^ and CD45^+^ cells. CD11b purity of the MACS-isolated samples was estimated as the percent of single cells positive for CD11b. CD11b^+^/CD45^high^ populations were manually gated blind to treatment condition. Outcome measures and findings across treatment groups were robust to the use of distinct antibodies and fluorochromes. Thus, all flow cytometry data within each treatment group were combined for subsequent presentation and analysis. An independent samples t-test was performed (Graph Pad Prism v9) to compare the proportion of CD11b^+^ cells contained in the CD45^high^ gate across treatment groups with alpha set to 0.05.

**Fig. 2.**
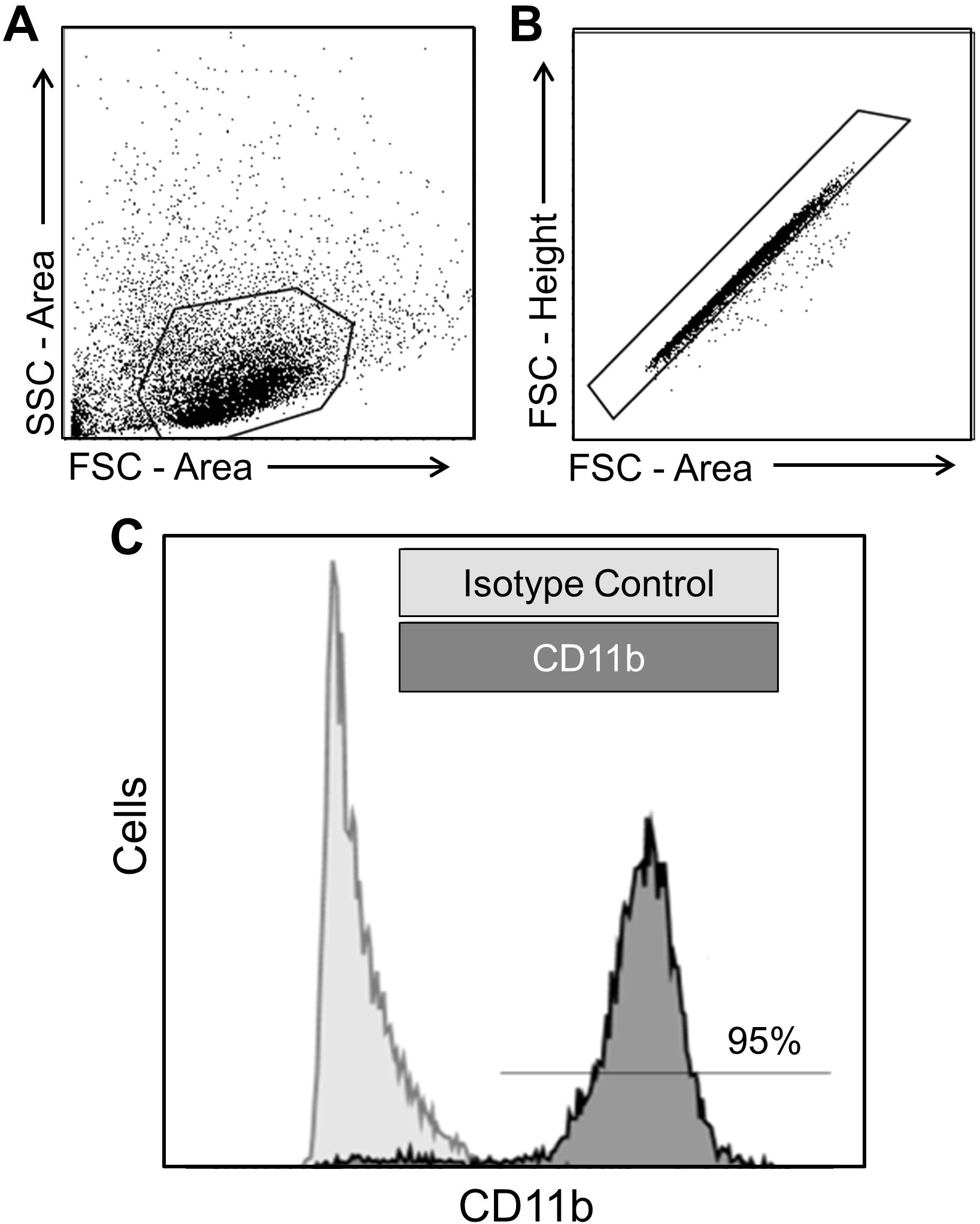
MACS-isolated CD11b^+^ cells show a high degree of purity. Gating strategies using forward scatter (FSC) and side scatter (SSC) profiles exclude (**A**) cell debris and (**B**) cell multiplets from the MACS-isolated CD11b^+^ cell suspension. (**C**) Representative data of flow cytometry. Relative to signal from isotype controls, ∼95% of the single cells show expression for CD11b.

## Results

### The isolated CD11b^+^ cells show a high degree of purity and viability

To assess the purity of the MACS-isolated CD11b**^+^** cells from the brains of aged APP/PS1mice, flow cytometry was used to quantify the cells with cell surface expression of CD11b (Fig. 2). We observed that 95% of the MACS-isolated cells expressed CD11b (Fig. 2C). The MACS-isolated CD11b**^+^** cells also showed a high degree of viability as assessed by trypan blue staining (95% viable). These findings indicate that our cell isolation method is effective, yielding highly pure and viable CD11b^+^ cells isolated from the brains of aged APP/PS1 mice.

### The vast majority of CD11b^+^ cells isolated from the brains of aged APP/PS1 mice have transcriptional profiles that align with microglia and myeloid cells

Next, scRNAseq was performed on the MACS-isolated CD11b^+^ cells from the brains of vehicle controls and SCF+G-CSF-treated old APP/PS1 mice. Gene transcription profiles were analyzed in a total of 19,008 cells, and tSNE plots were constructed to visualize expression patterns (Fig. 3). Unsupervised cluster analysis across all cells revealed 14 relatively unique clusters, which were color-coded and ranked according to cell count (Fig. 3A).

**Fig. 3.**
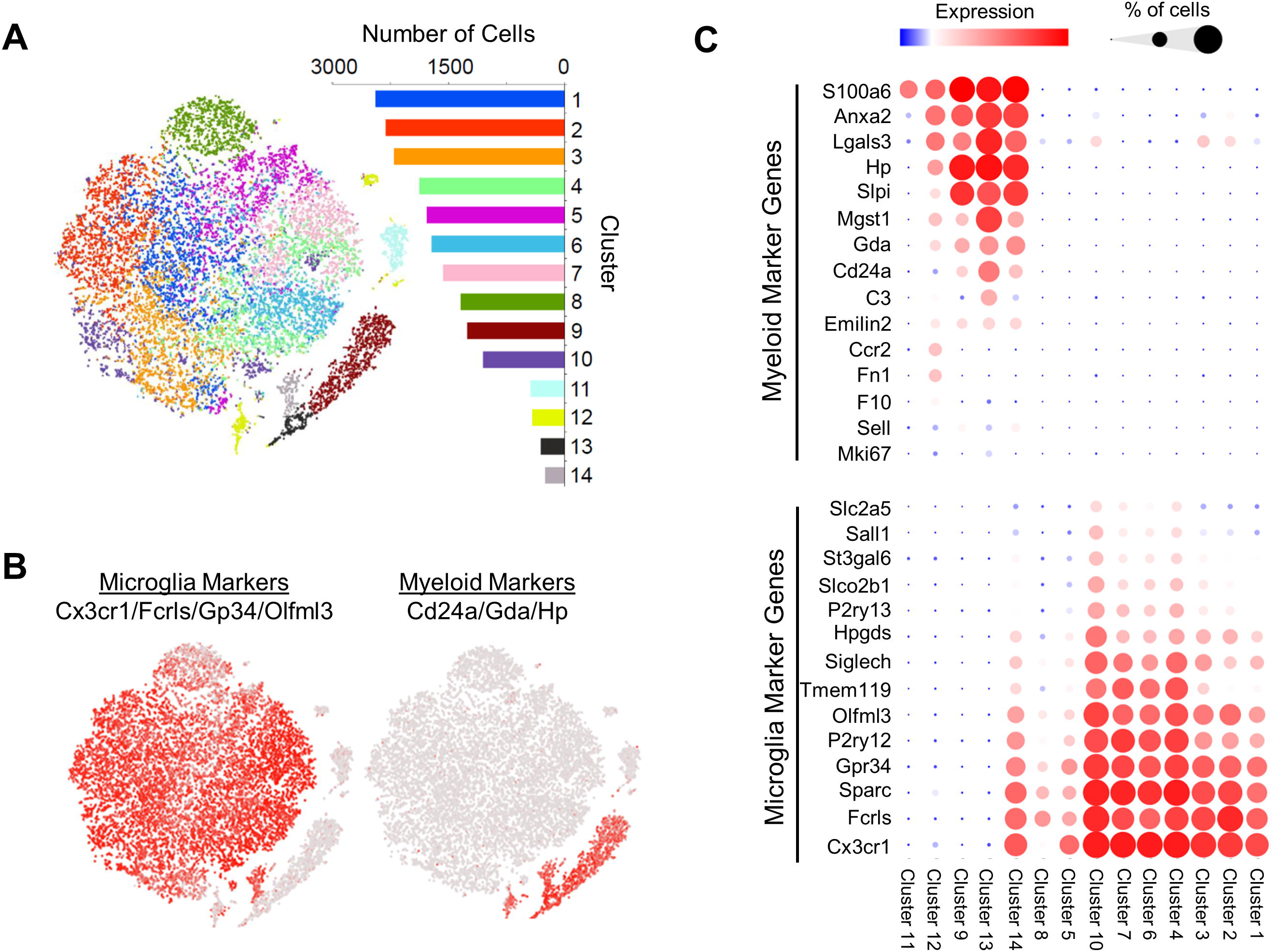
Cluster analysis of the single-cell RNA sequencing dataset reveals 14 cell clusters largely differentiated by marker genes of microglia and peripherally derived myeloid cells. (**A**) t-distributed stochastic neighbor embedding (tSNE) dimensionality reduction plots of 19,008 cells that are classified into 14 unique graph-based clusters defined and color-coded by unique transcriptional profiles. The number of cells in each cluster is shown in the corresponding bar graph. (**B**) Expression (in red) of microglial marker genes across tSNE plots indicates that the vast majority of clusters (1-8, 10, 14) associate with a microglia transcriptional signature. Expression (in red) of peripherally derived myeloid cell marker genes across tSNE plots indicates that several clusters (9, 12-14) associate with a myeloid cell transcriptional signature. Note, cluster 14 highly expresses genes associated with both cell classes. (**C**) Clusters 1-8, and 10 show enrichment of microglia gene profiles, while clusters 9, 12, and 13 show enrichment of myeloid cell profiles. Cluster 14 shows expression of marker genes of both cell types, while cluster 11 does not show expression of genes in either category. Bubble plots: the diameters of the circles correspond to the percentage of cells that express a given gene, while the color intensities of the circles correspond to the magnitude of expression.

The cell clusters highly expressed numerous genes commonly used to identify CD11b^+^ microglia and macrophages. Cst3, Lyz2, and Tyrobp showed relatively robust expression across individual clusters (Fig. S1) (Supplemental data File 1). In contrast, these clusters did not express marker genes of neurons, neuron progenitor cells, astrocytes, oligodendrocytes, oligodendrocyte progenitor cells, vascular cells, and fibroblasts (Fig. S1). These results confirm the purity of the MACS-isolated CD11b^+^ cells and suggest that the isolated CD11b^+^ cells are primarily composed of microglia and peripherally-born myeloid cell populations.

To further characterize the primary cell classes contained in each cluster, we analyzed the expression of gene sets identified by Haage and coworkers [79] to differentiate microglia from myeloid cells, thought to represent monocytes/macrophages. We found that the majority of clusters (clusters 1-8, and 10; containing 85.9% of all cells) aligned with microglia gene profiles (Cx3cr1^+^, Fcrls^+^, Sparc^+^, Gpr34^+^, P2ry12^+^, Olfml3^+^, Tmem119^+^, Siglech^+^, Hpgds^+^, P2ry13^+^, Slco2b1^+^, St3gal6^+^, Sall1^+^, and Scl2a5^+^) but not monocyte/macrophage profiles (S100a6^+^, Anxa2^+^, Lgals3, Hp^+^, Slpl^+^, Mgst1^+^, Gda^+^, CD24a^+^, C3^+^, Sell^+^, Emilin2^+^, Mki67^+^, F10^+^, Fn1^+^, and Ccr2^+^) (Fig. 3B-C; Fig. S2). In stark contrast, gene profiles of clusters 9, 12, and 13 (containing 11.8% of cells) largely aligned with those of monocytes/macrophages but not microglia (Fig. 3B-C). Consistent with these results, as compared to clusters 1-8 and 10, clusters 9, 12, and 13 showed significantly higher expression of Lgals3 (Fig. 3C; Fig. S2), an additional proposed marker gene of monocytes/macrophages following their migration to the brain [80]. Cluster 14 (containing 1.3% of cells) is a unique cluster showing robust expression of both microglial and monocyte/macrophage marker genes (Fig. 3C). Subsequent analysis of cluster 14 demonstrated significant expressions of S100a6, Anxa2, Hp, Lgals3, Slpi, and Gda (all monocyte/macrophage markers) in Cx3cr1^+^, P2ry12^+^, Fcrls^+^, Gpr34^+^, and Sparc^+^ cells (Table S1), suggesting robust co-expression of monocyte/macrophage and microglial gene sets in a small group of CD11b^+^ cells.

These collective results largely point to differentiated cell clusters of microglia and monocyte/macrophages. However, we note that given the chronic and severe neuropathological and neuroinflammatory conditions in the brains of aged APP/PS1 mice, it remains challenging to definitively distinguish microglia from peripherally derived macrophages based on transcriptional profiles alone [79,81]. Additionally, microglia and/or monocytes/macrophages express genes that are also expressed and/or used as marker genes in neutrophils [82–85], particularly under inflammatory conditions. These findings present a challenge to definitively differentiate sub-types of these cells. Considering these challenges, here we classify cell clusters 1-8, and 10 as microglial signature (MG-sig) clusters based on their microglial cell-like transcriptional profiles and clusters 9, 12, and 13 as myeloid signature (Mye-sig) clusters based on their transcriptional profiles that are similar to peripherally derived myeloid cells or monocytes/macrophages. Likewise, we classify cluster 14 as a MG/Mye-sig cluster based on a transcriptional profile that aligned with both microglia and monocytes/macrophages.

While cluster 11 (containing 2.3% of cells) showed high expression of S100a6, very little expression was observed of the remaining microglial and monocyte/macrophage gene markers (Fig. 3C). Inspection of gene sets associated with additional immune cell classes revealed relatively high expressions of Cd3d, Cd8b1, and Cd8a in cluster 11 (Fig. S1), demonstrating a profile consistent with that of T-cells [86].

Altogether, these findings suggest that the CD11b^+^ cells isolated from the brains of aged APP/PS1 mice have transcriptional profiles that align with those of microglia and peripheral myeloid cells, with a small population of cells showing a profile of T-cells. Given prior findings that SCF+G-CSF treatment induces changes in microglia and macrophages to engulf Aβ [22], the current study focuses strictly on the microglial cell-like MG-sig clusters and the myeloid cell-like Mye-sig clusters.

### Microglial signature clusters lie across a gradient of transcriptional profiles largely associated with activation and disease states

Differential gene analysis across cell clusters revealed gene sets relatively unique to each cluster. Across the MG-sig clusters, many top differentially expressed genes (DEG) were associated with homeostatic (e.g., Tmem119, P2ry12) or reactive disease states (e.g., Apoe, Cst7, Trem2, Itgax, Lpl, and Clec7a) [87, 88]. Emerging evidence demonstrates a role of highly reactive disease-associated microglia (DAM) [72] in slowing the progression of AD pathology [88]. Interestingly, this DAM state is associated with changes in expression levels of many of the top marker genes we noted across our MG-sig clusters. Moreover, DAM can take on distinct inflammatory profiles that correspond to phagocytic efficacies [89]. Given the overlap of DAM gene sets and our cluster-specific marker genes, we probed expression patterns of curated gene sets linked to homeostatic and DAM profiles [87], and to inflammation across clusters. To best characterize the diversity of the MG-sig clusters (1-8, and 10), genes with robust or variable expression patterns across clusters were selected and shown in Figure 4. While all MG-sig clusters expressed high levels of select DAM signature genes, including Trem2, Apoe, and Tyrobp, select homeostatic and DAM marker genes were variably expressed and generally demarcated the clusters (Fig. 4). Homeostatic genes (e.g., Tmem119, P2ry12, and Glul) were robustly enriched in clusters 4, 6, and 7, coinciding with a reduction of some DAM genes (e.g., Lpl, Fabp5, Clec7a, Cst7). Homeostatic genes, however, were downregulated in clusters 1, 2, 3, 5, and 8. Clusters 5 and 8 showed modest expression of lysosomal marker gene Cd68 and increased expression in relatively few variably expressed DAM genes (e.g., Cst7 and Fabp5). In contrast, clusters 1, 2, and 3 displayed heightened expression of Cd68 and the DAM genes Lyz2, Cst7, Clec7a, Fabp5, Spp1, Lpl, Itgax, Apoe, and Csf1, consistent with other reports suggesting an advanced or later-stage DAM profile [88] thought to further regulate phagocytic and lipid metabolic functions. Cluster 3 also expressed relatively high levels of Nfkbiz, Tnf, and Il1b genes, suggesting an inflammatory profile.

**Fig. 4.**
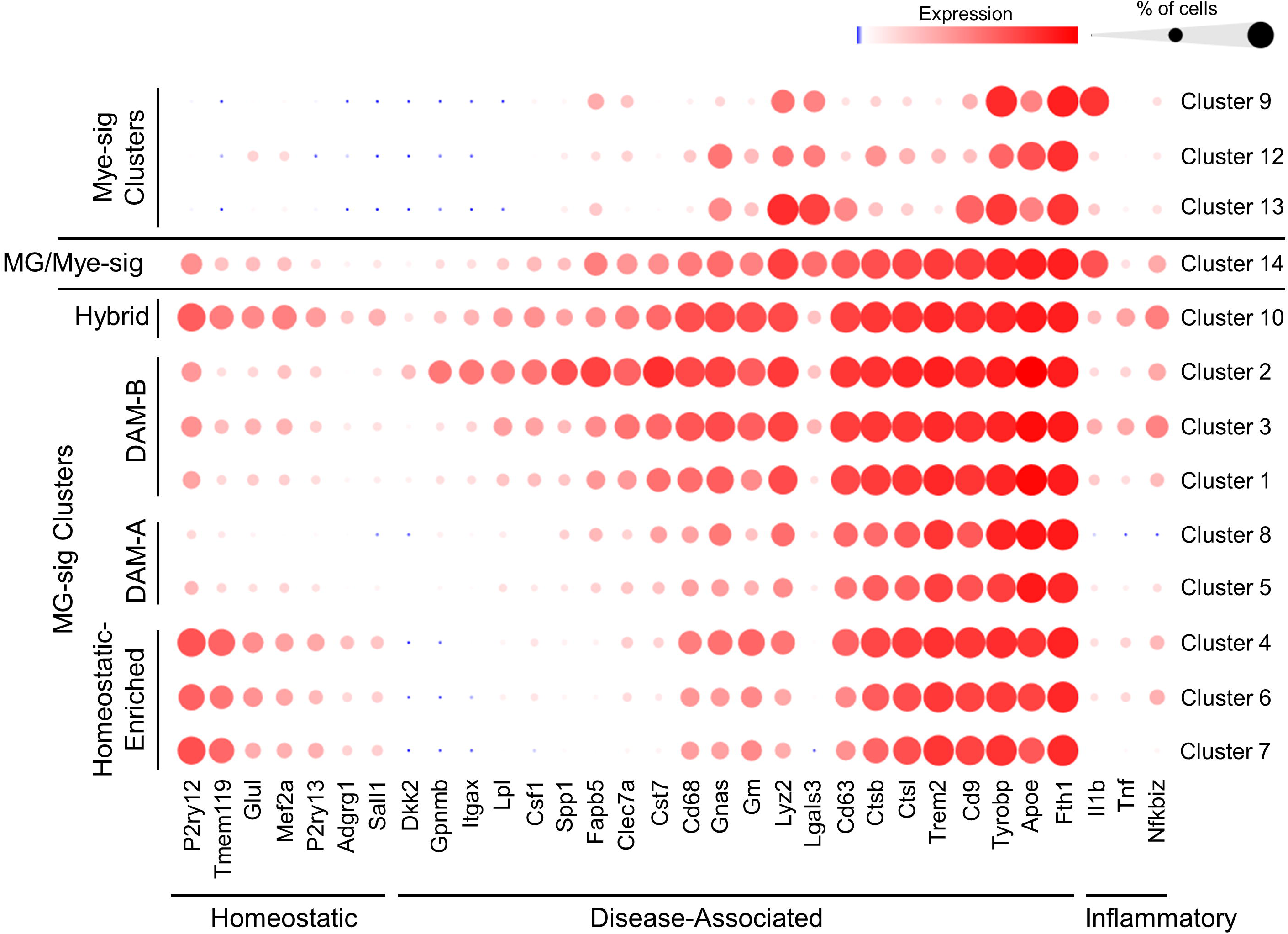
Expression profiles of homeostatic, DAM, and inflammatory gene sets reveal a gradient of activation states across clusters of CD11b^+^ cells isolated from the brains of aged APP/PS1 mice. Expression of select gene sets linked to homeostatic microglia, disease associated microglia (DAM), and inflammation are plotted across cell clusters. Homeostatic gene markers (e.g. Tmem119, P2ry12) are highly expressed in microglia-signature clusters 4, 6, and 7, while these clusters show relatively little expression of select DAM gene markers (e.g., Itgax, clec7a, Cst7, Lpl, Fabp5). These clusters were classified as MG-sig clusters with relative enrichment of homeostatic marker genes (i.e., homeostatic-enriched). In contrast, MG-sig cell clusters 1, 2, 3, 5, and 8 show high expression DAM genes with relatively little expression of homeostatic genes. These MG-sig clusters are classified as DAM-A (clusters 5 and 8) and DAM-B (clusters 1, 2, 3) according to the numbers of expressed DAM genes and levels of expression, with DAM-B showing a higher expression of more DAM genes. MG-sig cluster 10 has a transcriptional profile relatively enriched in both homeostatic and DAM genes and is characterized as a hybrid cluster. Mye-sig clusters 9, 12, and 13 while also expressing Tyrobp, Apoe, and Fth1 are largely associated with selective increases in Lgals3, and reduced expression of the majority of DAM genes. MG/Mye-sig cluster 14 shows a DAM-B profile. Bubble plots: the diameters of the circles correspond to the percentage of cells that express a given gene, while the color intensities of the circles correspond to the magnitude of expression.

MG-sig cluster 10 showed robust expression of the majority of DAM and homeostatic marker genes (Fig. 4). This profile was confirmed in individual Tmem119^+^ and P2ry12^+^ cells (Table S2), suggesting high levels of DAM and homeostatic marker genes in a subset of MG-sig cells. This cluster additionally showed relatively high expression levels of inflammatory genes (Nfkbiz, Tnf, and Il1b). These findings reveal a population of MG-sig cells characterized by high levels of homeostatic, DAM, and inflammatory signature genes.

The transcriptional profiles across MG-sig cells found here, appear to represent a wide range of transitional states, that, with the exception of a small subset (cluster 10), highlight an inverse relationship between enrichment of homeostatic genes and enrichment of DAM genes. For subsequent analysis, the MG-sig clusters were grouped into 4 distinct hubs based on their relative expression profiles of homeostatic and DAM gene sets (Fig. 4): 1) an MG-sig cluster hub showing relatively high expression of homeostatic genes and relatively low expression of DAM genes (clusters 4, 6, and 7; characterized as a homeostatic-enriched hub), 2) a relatively reactive MG-sig hub with modest enrichment of DAM signature genes (clusters 5 and 8; characterized as DAM-A), 3) an MG-sig hub relatively enriched with a high degree of DAM signature genes that corresponds to an advanced DAM stage (clusters 1, 2, and 3; characterized as DAM-B), and 4) an MG-sig hub highly enriched with both homeostatic and DAM genes (cluster 10; characterized as a hybrid hub). While we observed between-cluster and intra-cluster variation, our cluster hub classifications reflect robust changes in the transcriptional profiles of cluster-specific marker genes (Fig. 4). It is worth noting that all MG-sig clusters show some degree of expression of homeostatic and DAM gene sets. Our cluster hub classifications are relative based on changes in gene expression levels and changes in the percentage of cells that express related marker genes. For example, while the homeostatic-enriched cluster hub is defined based on relatively enriched expression of homeostatic genes and relatively low expression of some DAM genes, given the high expression of other core DAM genes, this does not necessarily mean that cells in the MG-sig homeostatic-enriched cluster hub are transcriptionally comparable to homeostatic microglia in the healthy brain of a young adult mouse.

Cluster 14, which transcriptionally aligned with both microglial and monocyte/macrophage profiles, highly expressed several genes associated with DAM-B microglia (e.g., Trem2, Cd68, Cst7, Clec7a, Fabp5). While Mye-sig clusters also expressed high levels of DAM genes, including Fth1, Tyrobp, and Apoe, the Mye-sig clusters displayed large increases in the monocyte/macrophage and DAM marker gene Lgals3 and showed little expression of the remaining DAM genes, including Trem2 (Fig. 4).

### SCF+G-CSF treatment increases the proportion of Mye-sig cells in the brains of aged APP/PS1 mice

Next, we sought to determine whether SCF+G-CSF treatment alters the proportions of MG-sig and Mye-sig cells in the brains of aged APP/PS1 mice. Figure 5A shows t-SNE plots for visualizing cell clusters in vehicle control and SCF+G-CSF treatment. In the vehicle controls, 98.7% of the cells were contained in the MG-sig clusters (clusters 1-8, 10), and 1.3% in the Mye-sig clusters (clusters 9, 12, 13) (Fig. 5B; Fig. S3A and B). Cell distributions in the SCF+G-CSF-treated mice were shifted with a significantly reduced percentage of cells in the MG-sig clusters (79.9% of cells; p < 0.0001) and a significantly increased percentage of cells in the Mye-sig clusters (20.1% of cells; p < 0.0001) as compared to those in the vehicle controls (Fig. 5B; Fig. S3A and B). Strikingly, cells from the SCF+G-CSF-treated mice accounted for 95.3% (range across clusters: 82% – 99%) of the cells contained in the Mye-sig clusters (9, 12, 13; Fig. S3B). In line with these findings, expression levels of all monocyte/macrophage gene markers were significantly upregulated by SCF+G-CSF treatment (Fig. 5C). We also observed that cluster 14 cells, which highly co-expressed microglial and monocyte/macrophage gene profiles, were almost exclusively (96.4%) from SCF+G-CSF-treated mice (Fig. 5D). Altogether, these findings suggest that SCF+G-CSF treatment may enhance the recruitment of myeloid cells into the brains of aged APP/PS1 mice.

**Fig. 5.**
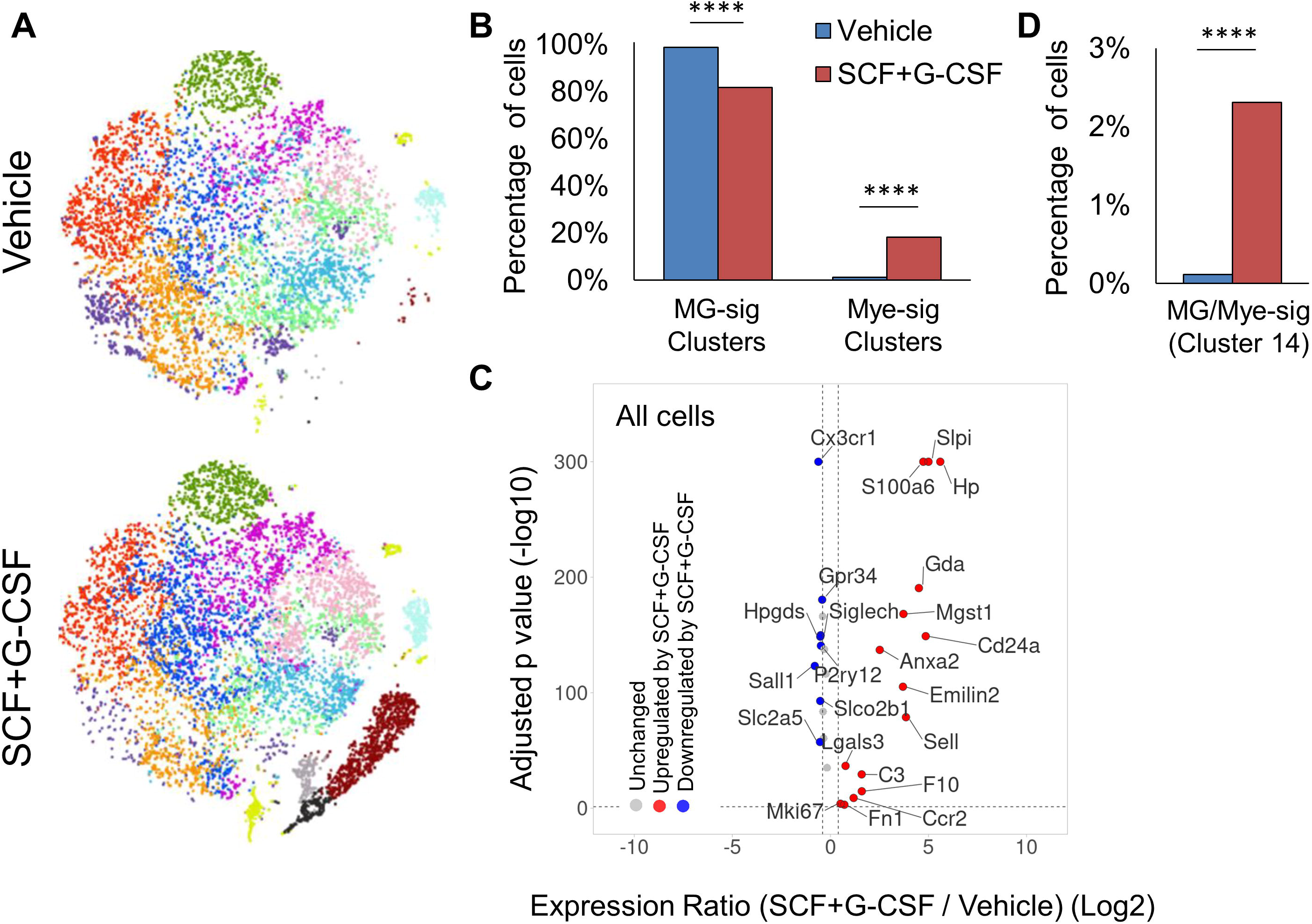
SCF+G-CSF treatment alters the distribution of cells across clusters, augmenting the prevalence of Mye-sig cells in the brains of aged APP/PS1 mice. (**A**) t-distributed stochastic neighbor embedding (tSNE) dimensionality reduction plots of cells separated by treatment. (**B**) SCF+G-CSF treatment shifts the distribution of CD11b^+^ cells with reduced proportions of MG-sig cells and increased proportions of Mye-sig cells in the brains of APP/PS1 mice. **** *p* < 0.0001. (**C**) A volcano plot highlights differential expression across SCF+G-CSF-treated and vehicle-treated groups of MG-sig and Mye-sig cell gene sets. Genes upregulated by SCF+G-CSF treatment are shown in red. Genes downregulated by SCF+G-CSF treatment are shown in blue. False discovery rate-corrected *p* values (-log10) are shown on the y-axis. The log2-transformed expression ratios are computed as expression levels in the SCF+G-CSF-treated mice compared to those in the vehicle control group and are plotted on the x-axis. Dotted lines indicate significance thresholds. (**D**) SCF+G-CSF treatment increases the number of MG/Mye-sig (cluster 14) cells. **** p < 0.0001.

To validate the scRNAseq data which suggest that SCF+G-CSF treatment increases the population of peripherally-born myeloid cells in the brain, we measured the cell surface expression of CD11b and CD45 using flow cytometry in separate aliquots of MACS-isolated CD11b^+^ cells. Previous studies show that the surface expression profiles of these proteins can be used to differentiate myeloid cells (CD11b^+^/CD45^high^) from microglia in the brain [90,91]. We found that SCF+G-CSF treatment significantly increased the proportion of CD11b^+^/CD45^high^ cells in the brains of aged APP/PS1 mice compared to the vehicle controls (p < 0.05; Fig. 6). These data support the scRNAseq findings suggesting that SCF+G-CSF treatment increases the population of peripherally derived myeloid cells or CD11b^+^/CD45^high^ active phagocytes in the brains of aged APP/PS1 mice.

**Fig. 6.**
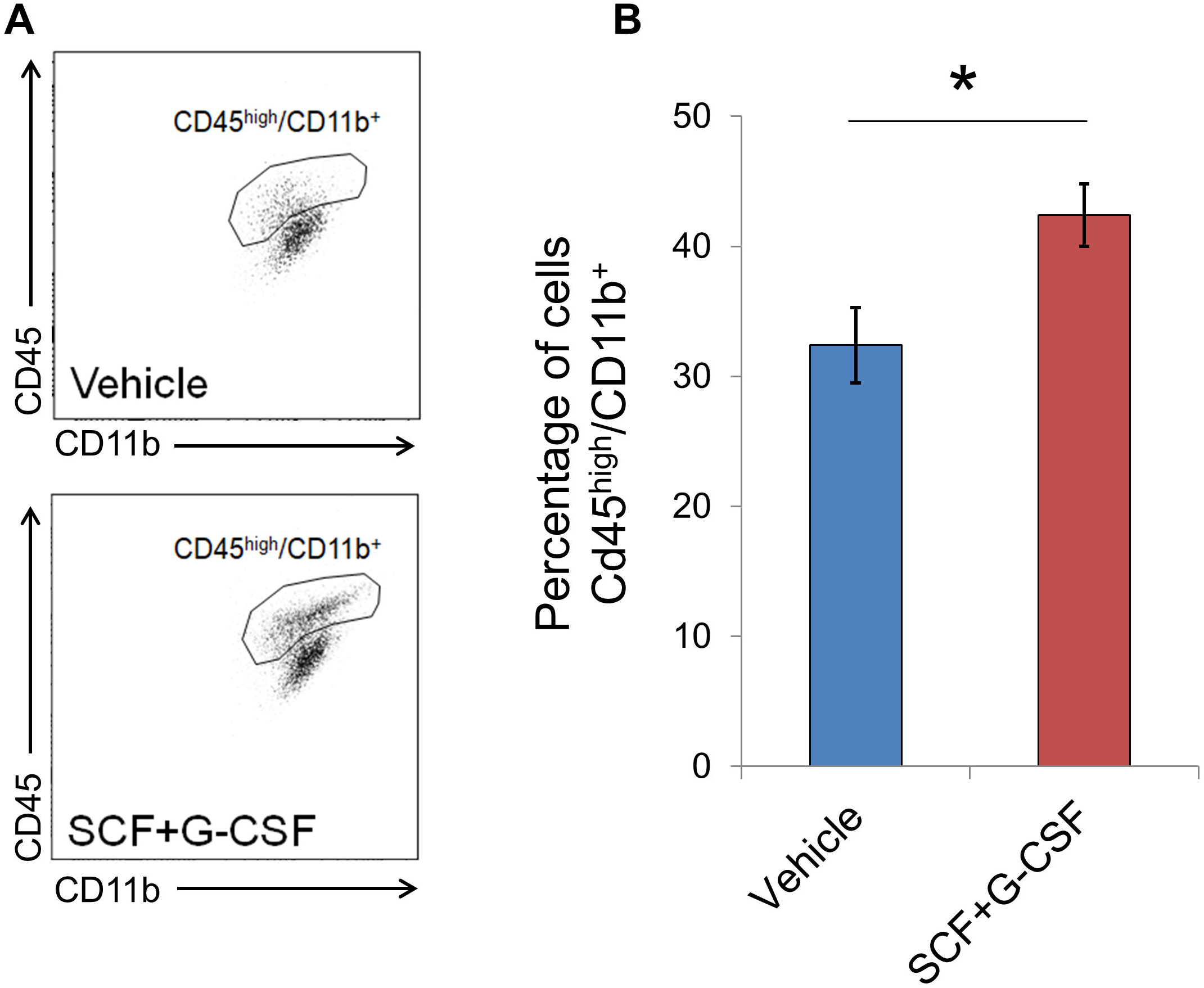
SCF+G-CSF treatment augments the recruitment of CD11b^+^/CD45^high^ cells into the brains of aged APP/PS1 mice. (**A**) Representative flow cytometry scatter plots show the population of CD45^high^/CD11b^+^ cells, thought to largely reflect peripherally derived myeloid cells/active phagocytes, in the brains of aged APP/PS1 mice treated with or without SCF+G-CSF. (**B**) SCF+G-CSF treatment increases the percentage of cells showing a CD45^high^/CD11b^+^ profile in the brains of aged APP/PS1 mice. Data are presented as mean ± SEM. * *p* < 0.05; t-test, n = 4 in each group.

### SCF+G-CSF treatment changes CD11b^+^ cell activation states in the brains of aged APP/PS1 mice

We next assessed the effects of SCF+G-CSF treatment in modifying the activation profiles in CD11b^+^ MG-sig clusters. SCF+G-CSF treatment significantly increased the percentage of MG-sig cells contained in the DAM-A hub (p < 0.0001, Fig. S4) and reduced the percentage of MG-sig cells in the hybrid cluster (i.e., cluster 10 that showed relatively high expressions of DAM, inflammatory, and homeostatic marker genes) (p < 0.0001, Fig. S4). The proportions of MG-sig cells contained in the homeostatic-enriched hub and DAM-B hub were not changed by SCF+G-CSF treatment (Fig. S4). These results suggest a shift away from inflammatory and homeostatic profiles in DAM-associated cells following SCF+G-CSF treatment. Consistent with these findings, differential expression analysis across all MG-sig clusters revealed that SCF+G-CSF treatment reduced the expression of inflammatory markers (e.g., Nfkbiz, Tnf, Il1b; Fig. S5A). However, SCF+G-CSF treatment also reduced the expressions of gene sets typically upregulated in DAM (e.g., Csf1, Axl, Itgax) as well as those typically downregulated in DAM (e.g., Egr1, Sall1, Jun; Fig. S5A), suggesting a variable or nuanced modification of transcriptional response by SCF+G-CSF treatment in DAM gene sets. These results were largely consistent upon comparison of gene sets across all clusters (Fig. S5B), with the notable exception of Il1b. Il1b was upregulated with SCF+G-CSF treatment (when pooling all clusters), an effect primarily driven by Il1b increases in the treatment-associated Mye-sig clusters (see Fig. 4).

### The most prominent transcriptome-wide responses to SCF+G-CSF treatment are largely comparable in MG-sig and Mye-sig clusters in the brains of aged APP/PS1 mice

Next, we performed transcriptome-wide differential gene analysis across experimental groups for unbiased identification of transcriptional changes in MG-sig clusters and Mye-sig clusters following SCF+G-CSF treatment. We first identified genes altered by SCF+G-CSF treatment when pooling all cell clusters together. We observed that the vast majority of differentially expressed genes (DEGs) was upregulated (n = 359 genes) by SCF+G-CSF treatment, rather than downregulated (n = 34 genes; Fig. S6). The top 10 DEGs ranked by significance and the number of differentially expressed transcripts (Fig. 7A) included four genes of the calcium-binding S100a family (S100a8, S100a9, S100a6, S100a11), the lipocalin family gene (Lcn2), the anti-inflammatory annexin A1 (Anxa1), the hemoglobin-binding haptoglobin (Hp), the anti-viral interferon-induced transmembrane gene (Ifitm1), the resistin-like molecule (Retnlg), and the whey acidic protein/four-disulfide core domain 21 (Wfdc21). These 10 genes were all upregulated by SCF+G-CSF treatment. This gene set collectively plays a prominent role in immune cell activation, such as microglia and macrophage-mediated inflammation, phagocytosis, and cell migration. Gene set enrichment analysis corroborated these findings linking the DEGs with immune responses, inflammatory responses, stress responses, and leukocyte migration (Fig. S7; Supplemental Data File 2).

**Fig. 7.**
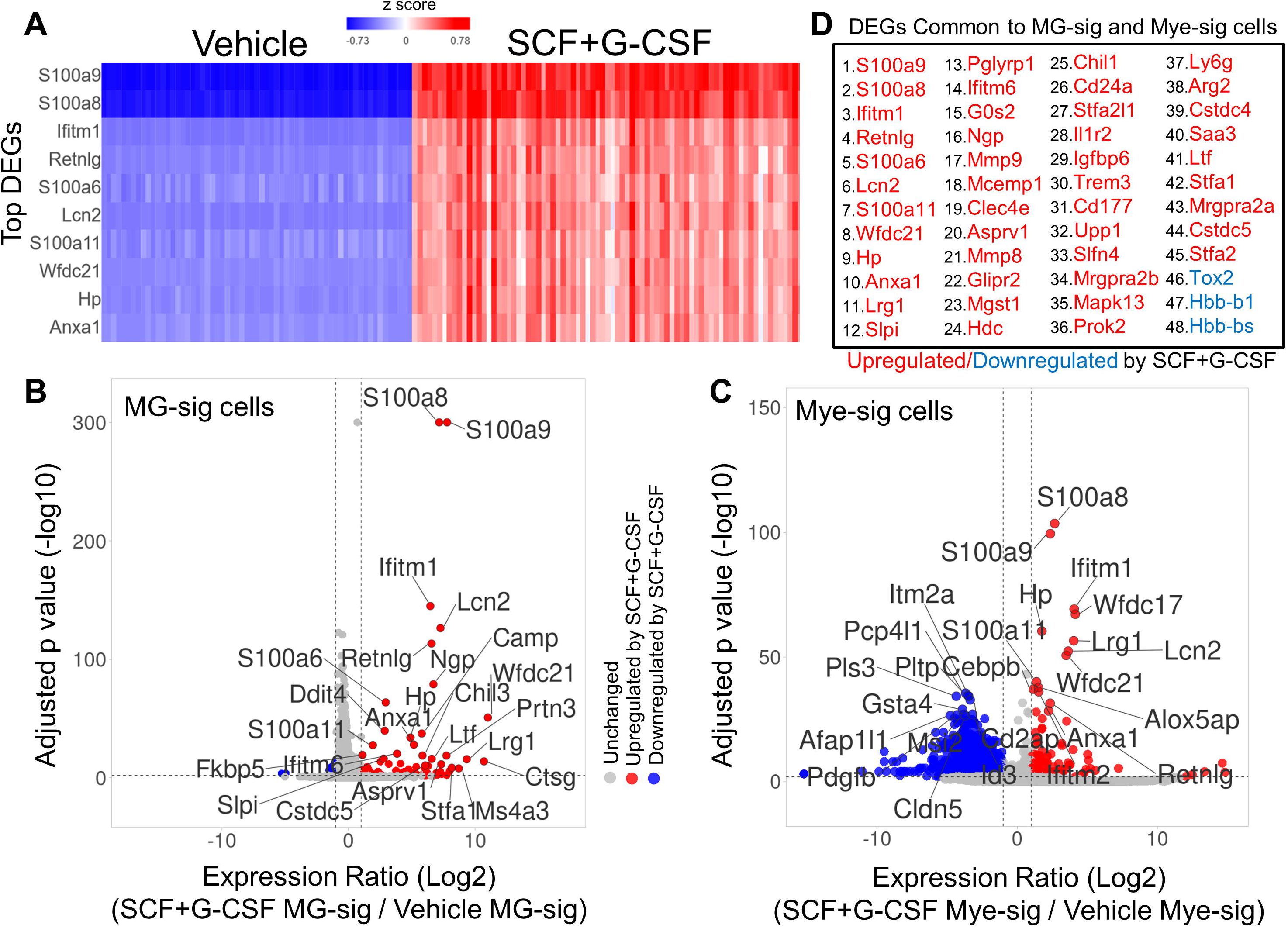
Transcriptome-wide responses to SCF+G-CSF treatment in MG-sig and Mye-sig cells in the brains of aged APP/PS1 mice. (**A**) A hierarchical clustering heat map illustrates the top 10 differentially expressed genes that are upregulated (in red) by SCF+G-CSF treatment across all cell clusters pooled together. The two upregulated genes S100a8 and S100a9 show the strongest effect following SCF+G-CSF treatment. **(B** and **C**) Volcano plots highlight genes differentially expressed by SCF+G-CSF treatment in MG-sig clusters (**B**) and in Mye-sig clusters (**C**). Genes upregulated by SCF+G-CSF treatment are shown in red. Genes downregulated by SCF+G-CSF treatment are shown in blue. In MG-sig cells, 70 genes are upregulated, and 18 genes are downregulated by SCF+G-CSF. In Mye-sig cells, 133 genes are upregulated, and 2637 genes are downregulated by SCF+G-CSF. False discovery rate-corrected *p* values (-log10) are shown on the y-axis. The log2-transformed expression ratios are computed as expression levels in SCF+G-CSF-treated mice relative to those in the vehicle control group. The log2-transformed expression ratios are plotted on the x-axis. Dotted lines indicate inclusion thresholds: ≥ 2-fold change, and FDR-corrected *p* value < 0.01. Note, *p* values are restricted to 300 decimal places. The top 25 genes differentially expressed are labeled. (**D**) A list of genes significantly altered (≥ 2-fold change, FDR-corrected *p* value < 0.01) by SCF+G-CSF treatment that are common to both MG-sig and Mye-sig cells is presented. Genes are ordered from lowest to highest FDR-corrected *p* value in the entire dataset.

The topmost genes upregulated by SCF+G-CSF treatment were largely comparable across MG-sig (Fig. 7B) and Mye-sig (Fig. 7C) clusters and across individual MG-sig cluster hubs (Fig. S8). These findings identify a stable transcriptional response following SCF+G-CSF treatment in brain immune cells of aged APP/PS1 mice. Expression profiles across individual cell clusters for the top 25 genes upregulated by SCF+G-CSF treatment are presented in Figure S9. Overlapping genes (n = 48) prominently altered by SCF+G-CSF treatment in both MG-sig and Mye-sig clusters (Supplemental Data File 3) are presented in Figure 7D.

While many of the DEGs upregulated by SCF+G-CSF treatment showed consistent profiles in MG-sig and Mye-sig clusters, we also noted distinct SCF+G-CSF-related transcriptional profiles across clusters. The majority of DEGs prominently altered by SCF+G-CSF treatment in MG-sig clusters were upregulated (n = 70 genes) rather than downregulated (n = 18 genes). By contrast, in Mye-sig clusters, 133 genes were upregulated, and 2637 genes were downregulated (Fig. 7B-C). These results suggest a more extensive transcriptional response to SCF+G-CSF treatment in the Mye-sig cells. Of note, the topmost genes downregulated by SCF+G-CSF in Mye-sig clusters included those associated with actin remodeling, cell adhesion and migration (e.g., Cd2ap, Pls3, Myo10, Afap1l1), vascular integrity (e.g., Pdgfb, Cldn5), and lymphocyte activation (e.g., Itm2a, Cd2ap).

### S100a8 and S100a9 are robustly expressed in MG-sig clusters and Mye-sig clusters in aged APP/PS1 mice following SCF+G-CSF treatment

Overall, the most striking effect following SCG+G-CSF treatment was upregulation of the genes S100a8 and S100a9. As S100a8 and S100a9 correlate with and interact directly with Aβ plaques, and they both are highly expressed in microglia and macrophages that surround plaques [92, 93], we further probed S100a8 and S100a9 expression profiles in our dataset. Violin density plots show the distributions of S100a8 and S100a9 expression in MG-sig and Mye-sig cells in vehicle control and SCF+G-CSF treatment groups (Fig. 8). The vast majority of the CD11b^+^ cells that highly expressed S100a8 or S100a9 were observed in the SCF+G-CSF-treated mice, with the highest expression in Mye-sig cells of SCF+G-CSF-treated mice.

**Fig. 8.**
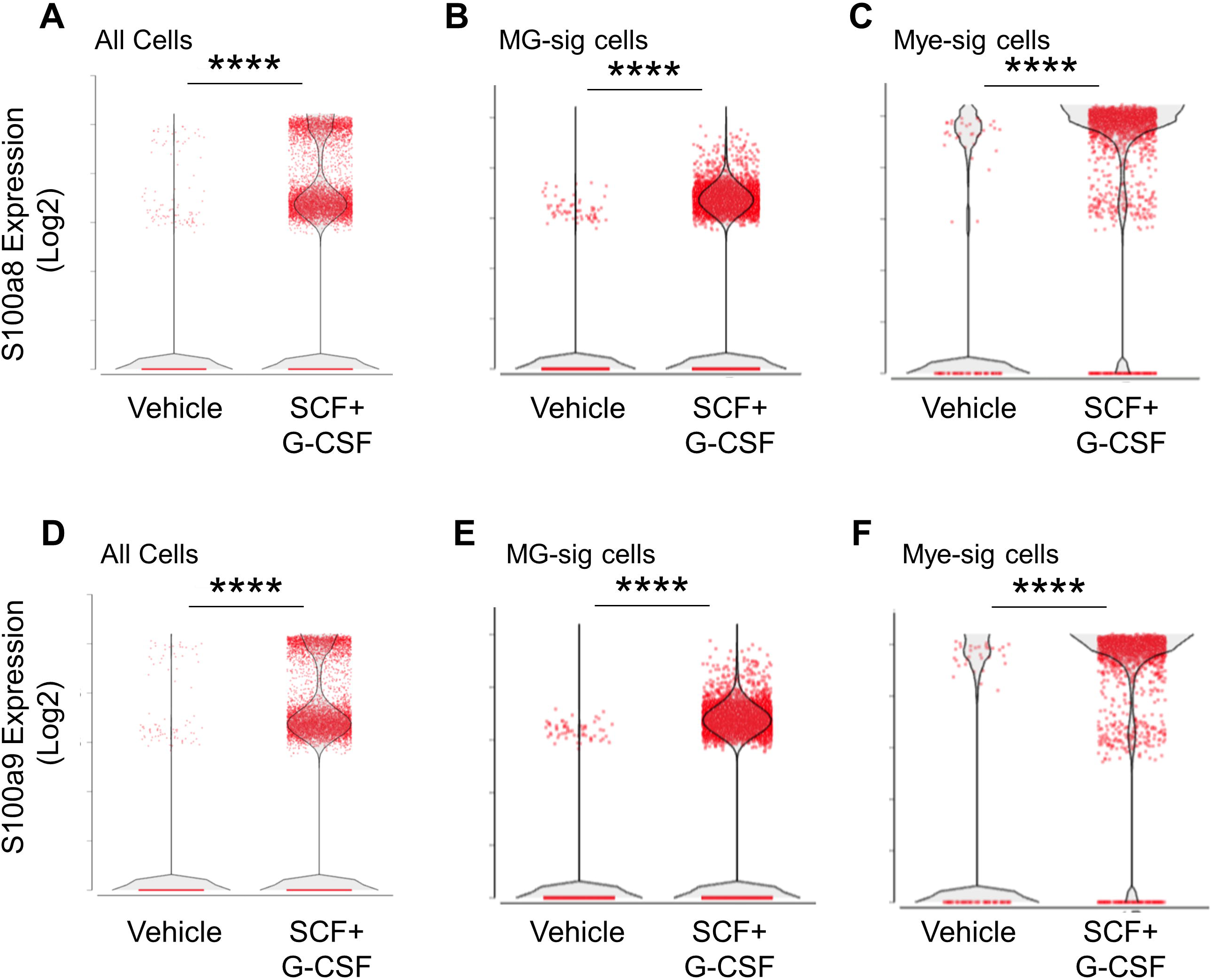
S100a8 and S100a9 are robustly upregulated by SCF+G+CSF treatment in aged APP/PS1 mice and most highly expressed in Mye-sig clusters. Violin density plots show that S100a8 (**A**-**C**) and S100a9 (**D**-**F**) are significantly upregulated by SCF+G-CSF treatment in cells pooled together across all clusters (**A** and **D**), in MG-sig clusters (**B** and **E**), and in Mye-sig clusters (**C** and **F**). Individual cells are shown as red dots. **** *p* < 0.0001.

To ensure that SCF+G-CSF treatment upregulated S100a8 and S100a9 (S100a8/9) expression in MG-sig cells rather than in small subsets of Mye-sig cells contained in the MG-sig clusters, we probed the expression levels of microglial gene markers in S100a8/9-positive cells and S100a8/9-negative cells. In MG-sig clusters, S100a8/9-positive cells co-expressed several specific gene markers of microglia, including Tmem119, at comparable levels to those in S100a8/9-negative cells (Fig. S10). This observation suggests that while expression of S100a8/9 is heightened in Mye-sig cells (see Fig. 8), the genes are also robustly enhanced in MG-sig cells following SCF+G-CSF treatment.

Interestingly, the topmost genes upregulated with SCF+G-CSF treatment in the entire dataset were co-expressed and upregulated in the S100a8/9-positive MG-sig cells compared to the S100a8/9-negative MG-sig cells as well as in the S100a8/9-positive Mye-sig cells compared to S100a8/9-negative Mye-sig cells (Fig. S11). These findings suggest that S100a8 and S100a9 are reliable marker genes associated with the prominent transcriptional responses to SCF+G-CSF treatment in brain immune cells in aged APP/PS1 mice.

### Network analysis of DEGs across experimental groups identifies highly inter-connected “hub” genes functionally connected to S100a8/9 expression

To identify potential interactions among the genes altered by SCF+G-CSF treatment, we constructed a functional network of gene-gene interactions among treatment-related DEGs (Fig. 9). A central hub within the network contained several of the top DEGs, including S100a8/9. Within this hub, we also noted highly inter-connected genes, including Il1b, Mmp9, Mpo, Cd44, and Rac2 (Fig. 9A and B). These findings suggest that central functions of CD11b^+^ cells changed by SCF+G-CSF treatment are associated with inflammation-like activation (Il1b, Mpo, Cd44), and cell motility and remodeling (Mmp9, Rac2, Cd44).

**Fig. 9.**
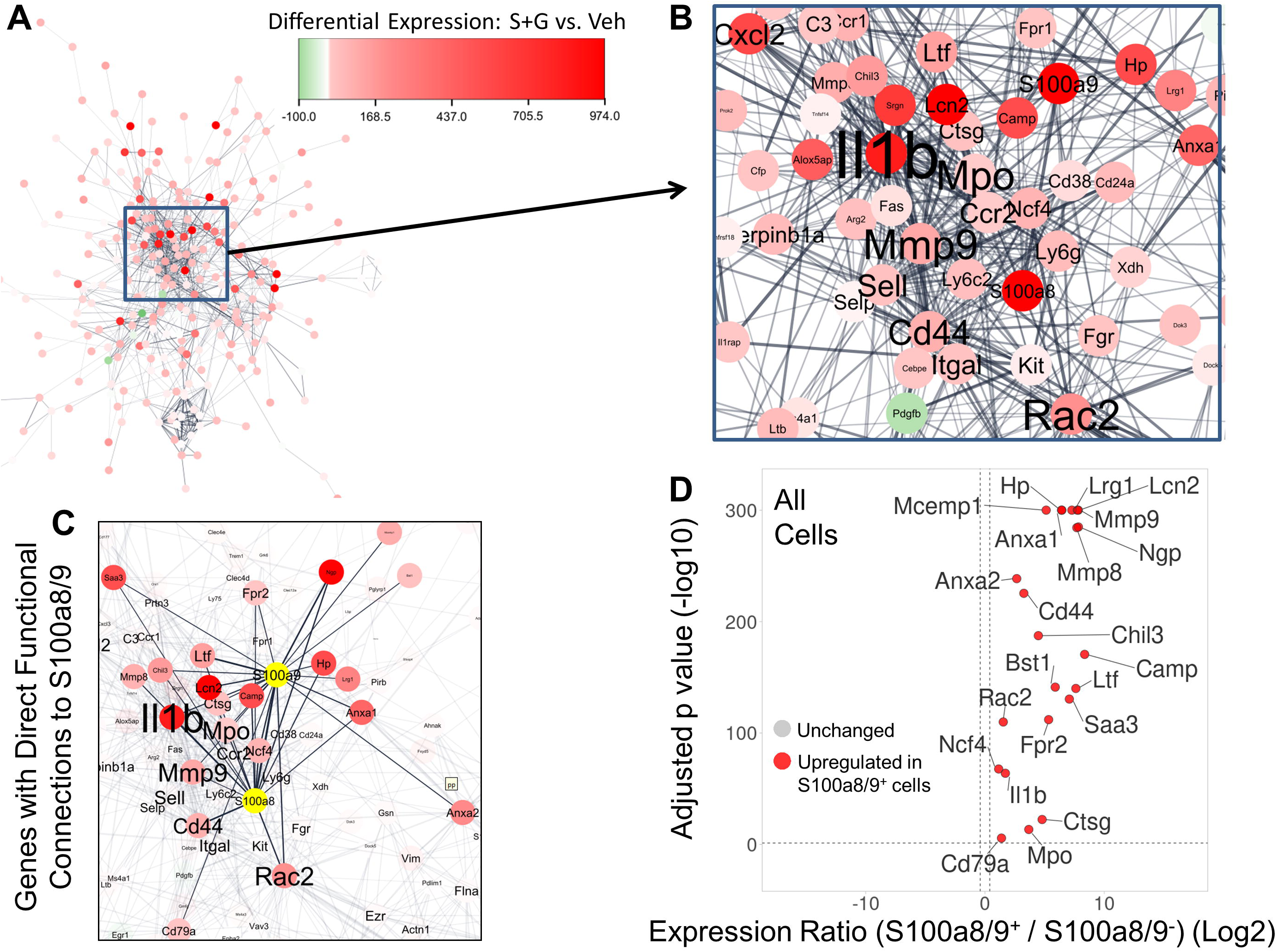
Functional network of genes differentially expressed by SCF+G-CSF treatment in aged APP/PS1 mice reveals highly inter-connected “hub” genes and those functionally linked with S100a8 and S100a9. (A) The genes significantly altered by SCF+G-CSF treatment (when all cells are pooled together) were used to create a functional gene-gene interaction network by querying their gene products in the STRING database. Each node corresponds to a gene significantly altered by SCF+G-CSF treatment. The edges (or links) between the nodes correspond to confirmed or potential functional connections. The color of the node corresponds to the intensity of differential expression, with red color indicating upregulation by SCF+G-CSF treatment. (B) A central hub of genes includes several of the top upregulated genes with SCF+G-CSF treatment (e.g., S100a8, S100a9, Anxa1, Lcn2, Hp). The size of the gene font corresponds to the extent of its connectivity in the network, identifying highly inter-connected “hub” genes, including Il1b, Mmp9, Rac2, and Cd44. (C) The 22 genes (Anxa1, Anxa2, Bst1, Camp, Cd44, Cd79a, Chil3, Ctsg, Fpr2, Hp, Il1b, Lcn2, Lrg1, Ltf, Mcemp1, Mmp8, Mmp9, Mpo, Ncf4, Ngp, Rac2, and Saa3) directly connected to S100a8 and S100a9 (S100a8/9) are highlighted. (D) This S100a8/9-linked gene set identified by network analysis is largely co-expressed and dramatically upregulated in individual S100a8/9-positive vs. S100a8/9-negative cells in the single-cell RNA sequencing dataset. Genes upregulated in S100a8/9-positive cells are shown in red. False discovery rate-corrected *p* values (-log10) are shown on the y-axis. The log2-transformed expression ratios are computed as expression levels in S100a8/9-positive cells relative to those in S100a8/9-negative cells and are plotted on the x-axis. Dotted lines indicate significance thresholds. Note, *p* values are restricted to 300 decimal places.

Twenty-two genes (Anxa1, Anxa2, Bst1, Camp, Cd44, Cd79a, Chil3, Ctsg, Fpr2, Hp, Il1b, Lcn2, Lrg1, Ltf, Mcemp1, Mmp8, Mmp9, Mpo, Ncf4, Ngp, Rac2, and Saa3) upregulated by SCF+G-CSF treatment were directly connected to S100a8/9 (Fig. 9C), suggesting direct functional interactions. Importantly, S100a8/9-positive MG-sig and Mye-sig cells showed enriched expressions of all 22 of these S100a8/9-connected genes (Fig. 9D; Fig. S12). We note that expression of the hub gene Il1b was unchanged in S100a8/9-positive MG-sig cells (Fig. S12A). However, expression of Il1b was upregulated by SCF+G-CSF treatment when all cells were pooled together (Fig. S5B) and in S100a8/9-positive Mye-sig cells (Fig. S12B). These findings demonstrate select contributions of S100a8/9-positive Mye-sig cells to SCF+G-CSF-related increases in the inflammatory hub gene Il1b.

Gene set enrichment analysis of S100a8/9-connected genes identified by network analysis showed enriched functions related to immune responses, inflammation (Cd44, Saa3, Anxa1, Il1b, Fpr2, Hp, Camp, S100a8, S100a9, Bst1, Mmp8), and leukocyte migration (Mmp9, Anxa1, Il1b, Rac2, Fpr2, Bst1, S100a8, S100a9) (Fig. S13). These findings are consistent with gene set enrichment analyses run on all DEGs (Fig. S7). Additionally, enriched Reactome pathways included degradation of the extracellular matrix (Mmp9, Mmp8, Ctsg, Cd44), implicating a mechanism related to cell motility and migration associated with SCF+G-CSF treatment. A complete list of enriched biological processes and Reactome pathways is provided in Supplemental Data File 2.

## Discussion

Our previous study demonstrated that SCF+G-SCF treatment activates microglia and macrophages to engulf and clear Aβ, an effect associated with reduced neuroinflammation and reconstructed neural connections in the brains of 26-27-month-old APP/PS1 mice [22]. In the present study, we employed a transcriptome-wide analysis of CD11b^+^ microglia and myeloid cells in the brains of 28-month-old APP/PS1 mice at the single-cell level following systemic treatment with SCF+G-CSF. Most notably, the findings of this current study show that SCF+G-CSF treatment 1) increases the proportions of immune cells in the brain that align with peripherally derived monocyte/macrophage transcriptional profiles, and 2) induces transcriptional responses in brain immune cells that are associated with cell activation, cell migration, inflammation, and Aβ clearance. These findings provide insight into how SCF+G-SCF treatment may modulate microglia and peripherally derived myeloid cells to reverse or mitigate neuropathology in the context of AD in old age.

In the current study, cluster analysis of MACS-isolated CD11b^+^ cells from the brains of aged APP/PS1 mice identified cell clusters with transcriptional profiles that aligned with microglial cell (MG-sig) and myeloid cell (Mye-sig) transcriptional profiles. The majority of these cell clusters (9 individual clusters: ∼86% of all cells) showed microglial cell-signature profiles. This observation is in line with previous studies showing that the vast majority of CD11b^+^ brain cells are microglia that comprise numerous transcriptionally-heterogeneous sub-populations in the aged, injured, or AD brain [94, 95, 96, 97, 98].

While all MG-sig clusters expressed high levels of signature DAM genes, including Trem2, Apoe, and Tyrobp, we found high cluster-to-cluster variability in the expression of homeostatic (e.g., Tmem119, P2ry12) and DAM-associated genes (e.g., Cst7, Trem2, Itgax, Lpl, and Clec7a). We also observed that between-cluster transcriptional variability in MG-sig cells was largely characterized by an inverse relationship between homeostatic and DAM profiles. This finding is well aligned with other reports showing transcriptional heterogeneity of microglia in AD that reflects transitionary stages from a relatively homeostatic to a highly reactive DAM phenotype [72, 98].

The findings of this present study also show that SCF+G-CSF treatment increases the proportion of DAM-like cells marked by relatively little homeostatic gene expression (most notably P2ry12). Homeostatic gene expression has been shown to inversely correlate with the proximity of microglia to Aβ plaques. A microglial phenotype characterized by reductions of P2ry12 has been found next to Aβ plaques in autopsy brain tissue from AD patients [99]. Moreover, our earlier study showed that P2ry12 expression is reduced in subsets of microglia that closely associate with Aβ plaques following SCF+G-CSF treatment [22]. Thus, our findings are consistent with an effect of SCF+G-CSF to augment a subset of DAM-like cells that closely surrounds Aβ plaques in the brains of aged APP/PS1 mice. Additionally, we observed that SCF+G-CSF treatment reduces pro-inflammatory gene markers (e.g., Il1b and Tnf) in MG-sig clusters. Reductions in pro-inflammatory genes, including Il1b, in DAM have been shown to be associated with enhanced phagocytosis of Aβ [89]. Taken together, the DAM transcriptional profiles reported here, increased by SCF+G-CSF treatment, may reflect increases in a subset of highly phagocytic but less inflammatory microglial cell-like cells that surround Aβ plaques. These effects may contribute to SCF+G-CSF-reduced Aβ burden and neuroinflammation in aged APP/PS1 mice [22]. Interestingly, similar effects are observed following granulocyte macrophage colony-stimulating factor (GM-CSF) treatment. GM-CSF treatment is shown to increase the number of reactive Iba1^+^ cells localized to Aβ plaques, an effect that corresponds to reductions in pro-inflammatory signaling and enhanced clearance of amyloid deposition [100, 101].

The findings of the present study reveal that SCF+G-CSF treatment dramatically increases the numbers of myeloid cells in the brains of aged APP/PS1 mice. The vast majority of CD11b^+^ cells contained in Mye-sig clusters are seen in the aged APP/PS1 mice treated with SCF+G-CSF, a finding consistent with the flow cytometry (immunoprofiling) results showing that SCF+G-CSF treatment increases CD11b^+^/CD45^high^ myeloid cells or active phagocytes [102] in the brains of aged APP/PS1 mice. We note that while these findings from transcriptomic and immunoprofiling experiments show the same directionality of effect (i.e., an increase in peripherally derived myeloid cells following SCF+G-CSF treatment), the percentage of cells with a CD11b^+^/CD45^high^ profile in flow cytometry experiments was higher than the percentage of cells in the Mye-sig clusters identified by transcriptomic profiling. It is possible that the distinct pre-processing methods across scRNAseq and flow cytometry experiments contribute to these differences. Additionally, it is possible that a subset of activated microglia that upregulate CD45 [81,102] could align with a microglial cell transcriptional signature yet a CD45^high^ immunophenotype.

Increased recruitment of bone marrow-derived macrophages in the brain following SCF+G-CSF treatment has been reported in middle-aged APP/PS1 mice [71]. Our findings extend the prior report by showing that SCF+G-CSF treatment enhances the recruitment of myeloid cells into the brain of APP/PS1 mice even in advanced age. Accumulating evidence suggests a prominent role of infiltrating myeloid cells in mitigating Aβ pathology in AD [31, 32, 33, 103,104]. GM-CSF treatment has been shown to increase monocyte migration [105], reduce Aβ plaques, and reverse cognitive deficits [100,101,106]. Consistent with these studies, our previous findings reveal that SCF+G-CSF treatment increases the association of bone marrow-derived macrophages with Aβ plaques and reduces Aβ deposition in APP/PS1 mice [71]. In addition, our unpublished two-photon live brain imaging study reveals an acute effect of SCF+G-CSF in reducing Aβ plaques during a 7-day treatment in APP/PS1 mice. Together with the observation of the current study showing SCF+G-CSF-increased CD11b^+^/CD45^high^ myeloid cells or active phagocytes [102] in the brains of aged APP/PS1 mice following a 5-day treatment of SCF+G-CSF, these findings point to a crucial role for peripherally derived myeloid cells in SCF+G-CSF-enhanced clearance of brain Aβ in the aged brain of APP/PS1 mice.

Transcriptome-wide analysis in the present study reveals dramatic and robust effects of SCF+G-CSF treatment in augmenting gene sets linked with increased immune responses, including inflammation and leukocyte migration. The top genes upregulated by SCF+G-CSF treatment (e.g., S100a8, S100a9, Ifitm1, Lcn2, Mmp9) are prominently associated with highly activated microglia and infiltrating myeloid cells, inflammation, and cell migration to sites of injury [92, 107–112]. Network analysis confirmed these effects by identifying hub genes that are functionally related to cell movement and remodeling (Mmp9, Rac2, Cd44), and inflammation (Il1b, Mpo, Cd44).

S100a8 and S100a9 were observed as the two most robustly upregulated genes following SCF+G-CSF treatment. Moreover, S100a8/9-positive microglia and peripherally derived myeloid cells highly co-express the topmost SCF+G-CSF-upregulated genes. These findings suggest that SCF+G-CSF treatment leads to a conserved or fundamental transcriptional profile characterized by upregulation of S100a8/9 in brain immune cells.

S100a family genes encode calcium-binding proteins that regulate fundamental intracellular processes, including intracellular calcium homeostasis, and cytoskeleton rearrangement. S100a8 and S100a9 are the most prominent calcium-binding proteins in bone marrow-derived phagocytic cells, and the secretion of S100a8 and S100a9 is associated with inflammatory processes [107, 113, 114]. Cytoskeletal remodeling during periods of inflammation appears intertwined with, and a pre-requisite of, cell migration. S100a9 activity is essential for trans-endothelial migration of leukocytes [107, 115]. S100a8/9 are linked to disease states, including AD. S100a8/9 are enriched in the brains and sera of AD patients and are thought to contribute to AD pathogenesis by promoting neuroinflammation and Aβ aggregation [92, 107, 116]. Thus, we are largely surprised to find that S100a8/9 are the topmost upregulated genes following SCF+G-CSF treatment which has therapeutic effects to mitigate Aβ plaque load and induce anti-inflammatory effects centrally in aged APP/PS1 mice [22].

However, relevant to our findings, inflammation and Aβ plaque load enhance S100a8/9 transcription and secretion by surrounding microglia and macrophages [92, 93]. Therefore, the finding that SCF+G-CSF treatment increases the numbers of activated microglia and macrophages that engulf Aβ plaques [22, 71] is consistent with SCF+G-CSF treatment-related enhancement of S100a8/9 transcription. Despite the prevailing view that S100a8/9 induces neuro-inflammation and contributes to AD pathogenesis, several lines of research point to S100a9-induced reductions in neural toxicity and inflammation caused by Aβ. Recent evidence demonstrates that while S100a9 accelerates Aβ plaque formation, it also infiltrates and compacts Aβ plaques and sequesters toxic soluble Aβ species, resulting in mitigation of neural toxicity [117]. Co-aggregation of S100a9 with Aβ also reduces both S100a9 inflammation and Aβ cytotoxicity [118], suggesting the inflammatory role of S100a9 is conditional on its molecular interactions or conformational state. Consistent with this interpretation, while secreted S100a8/9 can form a heterodimer to activate toll-like receptors to induce inflammation [107], high levels of extracellular calcium induce the formation of S100a8/9 heterotetramers, which may act through distinct pathways and reduce the inflammatory tone [119]. These findings are in line with anti-inflammatory properties of S100a8/9 under certain conditions, such as, combatting peripheral tissue damage during periods of heightened inflammation through sequestering the inflammatory cytokines Il1b and Tnf [107]. In the current study, SCF+G-CSF treatment induces downregulation of Il1b and Tnf and robust upregulation of S100a8/9 in MG-sig cells, suggesting a potential role for S100a8/9 in modulating inflammatory signaling in aged APP/PS1 mice.

While it remains possible that enhanced pro-inflammatory cytokines may contribute to phagocytic clearance of Aβ [23, 47, 48, 49, 50], inflammatory processes associated with S100a8/9 may be mitigated by anti-inflammatory-mediating genes co-upregulated by SCF+G-CSF treatment in microglia and peripherally-born myeloid cells. Several genes that are highly co-expressed in S100a8/9-positive MG-sig and Mye-sig cells or functionally linked with S100a8/9 expression (e.g., Anxa1, Anxa2, Chil3, Hp, Lrg1, Ngp, and Wfdc17) have been shown to confer anti-inflammatory or alternatively activated properties [120–123, 124, 125, 126, 127, 128] associated with tissue repair [39, 43, 129].

Moreover, several of the genes (Anxa1, Anxa2, Fpr2, S100a6, S100a11, Mmp9, Hp, Il1b) robustly upregulated with SCF+G-CSF treatment and linked with S100a8/9 are also associated with Aβ clearance and/or neuroprotection. These genes and their roles in AD and neuroprotection are briefly outlined below.

Annexins are calcium and phospholipid-binding proteins. Microglia Anxa1 mediates non-inflammatory phagocytosis of apoptotic cells. Under inflammatory conditions, Anxa1 treatment restores non-inflammatory clearance of apoptotic, but not healthy, neurons, and promotes the reduction of inflammatory processes in reactive microglia *in vitro* [128]. Anxa1 is upregulated in microglia that surround Aβ plaques in AD patients and AD mouse models [130]. Moreover, exogenous Anxa1 treatment *in vitro* enhances Aβ phagocytosis by microglia and reduces Aβ-stimulated transcription of pro-inflammatory genes, including Tnf in microglia [130]. In young 5xFAD mice, Anxa1 treatment reduces Tnf, mitigates blood-brain barrier pathology, and increases synaptic densities [131], demonstrating a neuroprotective role in AD pathology. The effects of Anxa1 in Aβ clearance are largely mediated by activation of its Fpr1/2 receptor [130]. Notably, our findings reveal that both Anxa1 and Fpr2 are also upregulated with SCF+G-CSF treatment. Moreover, an S100a11-encoded Anxa1 adapter protein enhances cytosolic and membrane-associated levels of Anxa1, which likewise confers neural protection in cerebral ischemia [132]. A similar anti-inflammatory and neuroprotective role of Anxa2 is reported in a TBI study [126].

S100a6 binds calcium and zinc and is found in cells that surround Aβ plaques. In a study targeting the contribution of zinc to Aβ aggregation, S100a6 treatment-reduced Aβ plaques in the brains of 22-month-old APP/PS1 mice were thought to reflect the ability of S100a6 to sequester excess zinc [133].

Hp encodes haptoglobin which is prominently identified as a hemoglobin-binding protein. The scavenging effects of haptoglobin in preventing neurotoxicity in the brain have been demonstrated in the condition of sub-arachnoid hemorrhage [134]. As cerebral micro-bleeds are associated with AD and correlate with Aβ burden [135], SCF+G-CSF-increased Hp expression may play a role in mitigating neuropathology in aged APP/PS1 mice. As shown in an *in vitro* study, haptoglobin associates with Aβ plaques and inhibits the formation of Aβ fibrils [136], suggesting direct functions of Hp to oppose Aβ deposition. High mobility group box-1 (HMGB1) is an activator of neuroinflammatory responses. HMGB1 inhibits microglial Aβ clearance, and factors that bind HMGB1 reduce Aβ load and suppress neuroinflammatory [137, 138]. Notably, it has been shown that haptoglobin also binds and sequesters HMGB1, which leads to reductions in tissue loss and pro-inflammatory TNF and increases in anti-inflammatory IL-10 in microglia and macrophages in a mouse model of stroke [127].

Mmp9 is a zinc and calcium-dependent endopeptidase. Through its proteolytic functions, Mmp9 plays a key role in the remodeling of the extracellular matrix, cell migration, and neuroplasticity [139]. Levels of Mmp9 are increased in AD, and Mmp9 shows the ability to degrade Aβ plaques [140, 141]. Moreover, neuronal over-expression of Mmp9 in 5xFAD mice shifts APP processing away from β- and γ-secretase pathways and toward an α-secretase pathway, which coincides with enhanced presynaptic densities [142]. An α-secretase pathway generates soluble APP-α, a non-amyloidogenic peptide that is reduced in AD and that promotes neuroprotection [143, 144].

Il1b, a classic pro-inflammatory cytokine, is associated with Aβ clearance. Overexpression of Il1b in the hippocampus of APP/PS1 mice reduces Aβ plaques, increases Iba1^+^ microglial cell and macrophage densities, and increases phagocytosis of Aβ [49, 50]. The findings of the present study show that SCF+G-CSF treatment increases Il1b expression in Mye-sig clusters, while SCF+G-CSF reduces Il1b in MG-sig clusters. It remains unclear if Mye-sig-derived Il1b upregulated by SCF+G-CSF treatment plays a prominent role to enhance phagocytic functions of CD11b^+^ myeloid cells to remove Aβ plaques.

Many of the genes upregulated by SCF+G-CSF treatment are activated by or modulate the concentrations of calcium, zinc, and/or iron (e.g., S100a8, S100a9, S100a11, S100a6, Lcn2, Hp, Mmp8, Mmp9, Anxa1, Anxa2, Fth1, Ltf). Indeed, metal sequestration and response to metal ions are enriched biological processes and Reactome pathways that are associated with SCF+G-CSF treatment in this study. As increased levels of metal ions play a causal role in AD pathogenesis [145, 146], the findings of this study support further research into metal sequestration functions following SCF+G-CSF treatment in the context of AD.

While future studies are needed to determine whether the gene sets outlined here play causal roles in resolving AD pathology in advanced age, we note that the robust transcriptional changes induced by SCF+G-CSF treatment remarkably overlap with those in CD11b^+^ cells in the brain during a period of functional recovery in a mouse model of TDP-43 proteinopathy [147]. It remains to be determined whether this gene set upregulated by SCF+G-CSF could serve as a reliable transcriptional profile of recovery and brain repair in the context of AD.

There are several limitations of this study. Microglial cell-like (MG-sig) and myeloid cell-like (Mye-sig) cell populations were distinguished in our dataset based on collated gene sets shown to differentiate the two cell classes [79]. If the severe neuropathology and neuroinflammation in the aged APP/PS1 mouse brain, and/or SCF+G-CSF treatment significantly altered the expression profiles of a high percentage of genes in our marker gene sets, our cell type classifications may be limited. It remains possible that the cell clusters defined herein represent mixed populations of microglial and myeloid cell types. Moreover, several myeloid cell classes have been shown to infiltrate the brains of transgenic mouse models of AD [148, 149]. While monocytes/macrophages are the primary bone marrow-derived myeloid cell population in the AD brain [31, 150], and these cells show a dramatic and selective influx to the brain following SCF+G-CSF treatment in middle-aged APP/PS1 mice [71], further work is needed to classify the populations of infiltrating myeloid cells following SCF+G-CSF treatment in the aged APP/PS1 brain. Likewise, use of tissue from a young, healthy, control mouse would better facilitate classification of distinct activations states (e.g., disease-associated and homeostatic). Indeed, corroboration of the transcriptional responses to SCF+G-CSF in identified subsets of cell classes using immunohistochemistry and/or single-molecule in situ hybridization will be useful to validate our findings.

In the current study we evaluated brain-wide transcriptional changes of CD11b^+^ cells following SCF+G-CSF treatment, precluding our ability to detect regional distinctions. As AD pathologies in the brain are spatially-diverse [151], it is possible that our study did not uncover SCF+G-CSF treatment effects limited to select brain regions.

We further stress that this scRNAseq study was run on pooled samples of three APP/PS1 mice treated with SCF+G-CSF and three APP/PS1 mice injected with vehicle solution. Thus, we were unable to distinguish the contributions of individual samples to the transcriptional profile associated with each condition, and consequently, could not determine individual variability across mice. While sample variability in individual transcriptional profiles could contribute to small changes across groups, large effects highlighted in our transcriptome-wide analysis were remarkably consistent in the vast majority of cluster hubs. Thus, individual animal variability did not likely confound or drive our primary findings.

In conclusion, the findings of this scRNAseq study reveal several potential cellular and molecular mechanisms by which systemic treatment of combined SCF+G-CSF modulates the functions of microglia and peripherally derived myeloid cells to clear and degrade Aβ plaques and mitigate AD pathology in the aged APP/PS1 mouse brain. Gene sets linked to immune cell activation, cell migration, and pro-inflammatory and anti-inflammatory processes are prominently enriched by the SCF+G-CSF treatment. Future studies will help clarify causal roles of the identified candidate mechanisms on brain repair following SCF+G-CSF treatment in age-related AD.

## List of abbreviations

Aβ: Amyloid-beta
AD: Alzheimer’s disease
DAM: Disease associated microglia
DEGs: Differentially expressed genes
D-PBS: Dulbecco’s phosphate-buffered saline
FDR: False discovery rate
G-CSF: Granulocyte colony-stimulating factor
MACS: Magnetic-activated cell sorting
MG-sig: Microglial transcriptional signature
Mye-sig: Peripherally-born myeloid transcriptional signature
scRNAseq: Single-cell RNA sequencing
SCF: Stem cell factor
TBI: Traumatic brain injury
tSNE: t-Distributed stochastic neighbor embedding
UMIs: Unique molecular identifiers

## Author Contributions

Conceptualization, L.R.Z.; methodology, R.S.G., M.K., and K.H.; formal analysis, R.S.G.; investigation, R.S.G.; resources, L.R.Z.; data curation, R.S.G.; writing—first draft preparation, R.S.G.; writing— review and editing, L.R.Z. R.S.G.; proofreading, M.K. and K.H.; visualization, R.S.G.; supervision, L.R.Z.; project administration and funding acquisition, L.R.Z. All authors read and approved the final version of the manuscript.

## Availability of data and materials

The datasets generated during the current study are available from the corresponding author on reasonable request.

## Declarations

### Ethics approval and consent to participate

This study was approved by Animal Care and Use Committee of SUNY Upstate Medical University.

### Consent for publication

Not applicable.

## References

1. Armstrong R. Risk factors for Alzheimer’s disease. Folia Neuropathol. 2019;57:87–105.

2. 2022 Alzheimer’s disease facts and figures. [cited 2022 Nov 19]; Available from: https://alz-journals.onlinelibrary.wiley.com/doi/10.1002/alz.12638

3. Rajan KB, Weuve J, Barnes LL, McAninch EA, Wilson RS, Evans DA. Population estimate of people with clinical Alzheimer’s disease and mild cognitive impairment in the United States (2020-2060). Alzheimers Dement J Alzheimers Assoc. 2021;17:1966–75.

4. Goedert M. Alzheimer’s and Parkinson’s diseases: The prion concept in relation to assembled Aβ, tau, and α-synuclein. Science. 2015;349:1255555.

5. Tiwari S, Atluri V, Kaushik A, Yndart A, Nair M. Alzheimer’s disease: pathogenesis, diagnostics, and therapeutics. Int J Nanomedicine. 2019;14:5541–54.

6. O’Brien RJ, Wong PC. Amyloid Precursor Protein Processing and Alzheimer’s Disease. Annu Rev Neurosci. 2011;34:185–204.

7. Seeman P, Seeman N. Alzheimer’s disease: β-amyloid plaque formation in human brain. Synap N Y N. 2011;65:1289–97.

8. Fein JA, Sokolow S, Miller CA, Vinters HV, Yang F, Cole GM, et al. Co-localization of amyloid beta and tau pathology in Alzheimer’s disease synaptosomes. Am J Pathol. 2008;172:1683–92.

9. Hanseeuw BJ, Betensky RA, Jacobs HIL, Schultz AP, Sepulcre J, Becker JA, et al. Association of Amyloid and Tau With Cognition in Preclinical Alzheimer Disease: A Longitudinal Study. JAMA Neurol. 2019;76:915–24.

10. Heneka MT, Carson MJ, El Khoury J, Landreth GE, Brosseron F, Feinstein DL, et al. Neuroinflammation in Alzheimer’s disease. Lancet Neurol. 2015;14:388–405.

11. Mormino EC, Kluth JT, Madison CM, Rabinovici GD, Baker SL, Miller BL, et al. Episodic memory loss is related to hippocampal-mediated beta-amyloid deposition in elderly subjects. Brain J Neurol. 2009;132:1310– 23.

12. Sato C, Barthélemy NR, Mawuenyega KG, Patterson BW, Gordon BA, Jockel-Balsarotti J, et al. Tau Kinetics in Neurons and the Human Central Nervous System. Neuron. 2018;97:1284–1298.e7.

13. Villemagne VL, Burnham S, Bourgeat P, Brown B, Ellis KA, Salvado O, et al. Amyloid β deposition, neurodegeneration, and cognitive decline in sporadic Alzheimer’s disease: a prospective cohort study. Lancet Neurol. 2013;12:357–67.

14. Bloom GS. Amyloid-β and Tau: The Trigger and Bullet in Alzheimer Disease Pathogenesis. JAMA Neurol. 2014;71:505–8.

15. Hurtado DE, Molina-Porcel L, Iba M, Aboagye AK, Paul SM, Trojanowski JQ, et al. A{beta} accelerates the spatiotemporal progression of tau pathology and augments tau amyloidosis in an Alzheimer mouse model. Am J Pathol. 2010;177:1977–88.

16. Lewis J, Dickson DW, Lin WL, Chisholm L, Corral A, Jones G, et al. Enhanced neurofibrillary degeneration in transgenic mice expressing mutant tau and APP. Science. 2001;293:1487–91.

17. Götz J, Chen F, van Dorpe J, Nitsch RM. Formation of neurofibrillary tangles in P301l tau transgenic mice induced by Abeta 42 fibrils. Science. 2001;293:1491–5.

18. Diaz A, Limon D, Chávez R, Zenteno E, Guevara J. Aβ25-35 injection into the temporal cortex induces chronic inflammation that contributes to neurodegeneration and spatial memory impairment in rats. J Alzheimers Dis JAD. 2012;30:505–22.

19. Faucher P, Mons N, Micheau J, Louis C, Beracochea DJ. Hippocampal Injections of Oligomeric Amyloid β- peptide (1–42) Induce Selective Working Memory Deficits and Long-lasting Alterations of ERK Signaling Pathway. Front Aging Neurosci. 2016;7:245.

20. Henriques A, Jégo P, Farrugia C, Callizot N. Intra-hippocampal injections of Aβ oligomers induce cognitive impairments associated with neurodegeneration and activation of microglia in senescent mice: Characterization of a novel animal model of Alzheimer’s disease. Alzheimers Dement. 2021;17:e054250.

21. Cai Z, Hussain MD, Yan L-J. Microglia, neuroinflammation, and beta-amyloid protein in Alzheimer’s disease. Int J Neurosci. 2014;124:307–21.

22. Guo X, Liu Y, Morgan D, Zhao L-R. Reparative Effects of Stem Cell Factor and Granulocyte Colony-Stimulating Factor in Aged APP/PS1 Mice. Aging Dis. 2020;11:1423–43.

23. He Z, Yang Y, Xing Z, Zuo Z, Wang R, Gu H, et al. Intraperitoneal injection of IFN-γ restores microglial autophagy, promotes amyloid-β clearance and improves cognition in APP/PS1 mice. Cell Death Dis. 2020;11:440.

24. Tian D-Y, Cheng Y, Zhuang Z-Q, He C-Y, Pan Q-G, Tang M-Z, et al. Physiological clearance of amyloid-beta by the kidney and its therapeutic potential for Alzheimer’s disease. Mol Psychiatry. 2021;26:6074–82.

25. Briggs R, Kennelly SP, O’Neill D. Drug treatments in Alzheimer’s disease. Clin Med. 2016;16:247–53.

26. Haeberlein S, Aisen PS, Barkhof F, Chalkias S, Chen T, Cohen S, et al. Two Randomized Phase 3 Studies of Aducanumab in Early Alzheimer’s Disease. J Prev Alzheimers Dis. 2022;9:197–210.

27. Liu KY, Villain N, Ayton S, Ackley SF, Planche V, Howard R, et al. Key questions for the evaluation of antiamyloid immunotherapies for Alzheimer’s disease. Brain Commun. 2023;5:fcad175.

28. Cummings J, Osse AML, Cammann D, Powell J, Chen J. Anti-Amyloid Monoclonal Antibodies for the Treatment of Alzheimer’s Disease. BioDrugs. 2024;38:5–22.

29. Lee CYD, Landreth GE. The role of microglia in amyloid clearance from the AD brain. J Neural Transm Vienna Austria 1996. 2010;117:949–60.

30. Kettenmann H, Hanisch U-K, Noda M, Verkhratsky A. Physiology of microglia. Physiol Rev. 2011;91:461–553.

31. Simard AR, Soulet D, Gowing G, Julien J-P, Rivest S. Bone marrow-derived microglia play a critical role in restricting senile plaque formation in Alzheimer’s disease. Neuron. 2006;49:489–502.

32. Town T, Laouar Y, Pittenger C, Mori T, Szekely CA, Tan J, et al. Blocking TGF-beta-Smad2/3 innate immune signaling mitigates Alzheimer-like pathology. Nat Med. 2008;14:681–7.

33. Yong VW, Rivest S. Taking Advantage of the Systemic Immune System to Cure Brain Diseases. Neuron. 2009;64:55–60.

34. Zhao Y, Wu X, Li X, Jiang L-L, Gui X, Liu Y, et al. TREM2 is a receptor for β-amyloid which mediates microglial function. Neuron. 2018;97:1023–1031.e7.

35. Fan Y, Ma Y, Huang W, Cheng X, Gao N, Li G, et al. Up-regulation of TREM2 accelerates the reduction of amyloid deposits and promotes neuronal regeneration in the hippocampus of amyloid beta1-42 injected mice. J Chem Neuroanat. 2019;97:71–9.

36. Wang Y, Cella M, Mallinson K, Ulrich JD, Young KL, Robinette ML, et al. TREM2 lipid sensing sustains the microglial response in an Alzheimer’s disease model. Cell. 2015;160:1061–71.

37. Karch CM, Goate AM. Alzheimer’s disease risk genes and mechanisms of disease pathogenesis. Biol Psychiatry. 2015;77:43–51.

38. Katsumoto A, Takeuchi H, Takahashi K, Tanaka F. Microglia in Alzheimer’s Disease: Risk Factors and Inflammation. Front Neurol. 2018;9:978.

39. Guo S, Wang H, Yin Y. Microglia Polarization From M1 to M2 in Neurodegenerative Diseases. Front Aging Neurosci. 2022;14:815347.

40. Shabab T, Khanabdali R, Moghadamtousi SZ, Kadir HA, Mohan G. Neuroinflammation pathways: a general review. Int J Neurosci. 2017;127:624–33.

41. Cherry JD, Olschowka JA, O’Banion MK. Neuroinflammation and M2 microglia: the good, the bad, and the inflamed. J Neuroinflammation. 2014;11:98.

42. Koenigsknecht-Talboo J, Landreth GE. Microglial Phagocytosis Induced by Fibrillar β-Amyloid and IgGs Are Differentially Regulated by Proinflammatory Cytokines. J Neurosci. 2005;25:8240–9.

43. Zhang Y, He M-L. Deferoxamine enhances alternative activation of microglia and inhibits amyloid beta deposits in APP/PS1 mice. Brain Res. 2017;1677:86–92.

44. Saresella M, Marventano I, Calabrese E, Piancone F, Rainone V, Gatti A, et al. A complex proinflammatory role for peripheral monocytes in Alzheimer’s disease. J Alzheimers Dis JAD. 2014;38:403–13.

45. Jimenez S, Baglietto-Vargas D, Caballero C, Moreno-Gonzalez I, Torres M, Sanchez-Varo R, et al. Inflammatory response in the hippocampus of PS1M146L/APP751SL mouse model of Alzheimer’s disease: age-dependent switch in the microglial phenotype from alternative to classic. J Neurosci Off J Soc Neurosci. 2008;28:11650–61.

46. Varnum MM, Ikezu T. The classification of microglial activation phenotypes on neurodegeneration and regeneration in Alzheimer’s disease brain. Arch Immunol Ther Exp (Warsz). 2012;60:251–66.

47. Herber DL, Mercer M, Roth LM, Symmonds K, Maloney J, Wilson N, et al. Microglial activation is required for Abeta clearance after intracranial injection of lipopolysaccharide in APP transgenic mice. J Neuroimmune Pharmacol Off J Soc NeuroImmune Pharmacol. 2007;2:222–31.

48. Majumdar A, Cruz D, Asamoah N, Buxbaum A, Sohar I, Lobel P, et al. Activation of microglia acidifies lysosomes and leads to degradation of Alzheimer amyloid fibrils. Mol Biol Cell. 2007;18:1490–6.

49. Rivera-Escalera F, Pinney JJ, Owlett L, Ahmed H, Thakar J, Olschowka JA, et al. IL-1β-driven amyloid plaque clearance is associated with an expansion of transcriptionally reprogrammed microglia. J Neuroinflammation. 2019;16:261.

50. Shaftel SS, Kyrkanides S, Olschowka JA, Miller JH, Johnson RE, O’Banion MK. Sustained hippocampal IL-1β overexpression mediates chronic neuroinflammation and ameliorates Alzheimer plaque pathology. J Clin Invest. 2007;117:1595–604.

51. Venegas C, Kumar S, Franklin BS, Dierkes T, Brinkschulte R, Tejera D, et al. Microglia-derived ASC specks cross-seed amyloid-β in Alzheimer’s disease. Nature. 2017;552:355–61.

52. Ising C, Heneka MT. Functional and structural damage of neurons by innate immune mechanisms during neurodegeneration. Cell Death Dis. 2018;9:1–8.

53. Briddell RA, Hartley CA, Smith KA, McNiece IK. Recombinant Rat Stem Cell Factor Synergizes With Recombinant Human Granulocyte Colony-Stimulating Factor In Vivo in Mice to Mobilize Peripheral Blood Progenitor Cells That Have Enhanced Repopulating Potential. Blood. 1993;82:1720–3.

54. Hess DA, Levac KD, Karanu FN, Rosu-Myles M, White MJ, Gallacher L, et al. Functional analysis of human hematopoietic repopulating cells mobilized with granulocyte colony-stimulating factor alone versus granulocyte colony-stimulating factor in combination with stem cell factor. Blood. 2002;100:869–78.

55. Welte K, Platzer E, Lu L, Gabrilove JL, Levi E, Mertelsmann R, et al. Purification and biochemical characterization of human pluripotent hematopoietic colony-stimulating factor. Proc Natl Acad Sci U S A. 1985;82:1526–30.

56. Zsebo KM, Wypych J, McNiece IK, Lu HS, Smith KA, Karkare SB, et al. Identification, purification, and biological characterization of hematopoietic stem cell factor from buffalo rat liver-conditioned medium. Cell. 1990;63:195–201.

57. Ping S, Qiu X, Kyle M, Hughes K, Longo J, Zhao L-R. Stem cell factor and granulocyte colony-stimulating factor promote brain repair and improve cognitive function through VEGF-A in a mouse model of CADASIL. Neurobiol Dis. 2019;132:104561.

58. Rahi V, Jamwal S, Kumar P. Neuroprotection through G-CSF: recent advances and future viewpoints. Pharmacol Rep PR. 2021;73:372–85.

59. Zhang SC, Fedoroff S. Cellular localization of stem cell factor and c-kit receptor in the mouse nervous system. J Neurosci Res. 1997;47:1–15.

60. Cui L, Murikinati SR, Wang D, Zhang X, Duan W-M, Zhao L-R. Reestablishing neuronal networks in the aged brain by stem cell factor and granulocyte-colony stimulating factor in a mouse model of chronic stroke. PloS One. 2013;8:e64684.

61. Qiu X, Ping S, Kyle M, Chin L, Zhao L-R. Long-term beneficial effects of hematopoietic growth factors on brain repair in the chronic phase of severe traumatic brain injury. Exp Neurol. 2020;330:113335.

62. Ping S, Qiu X, Gonzalez-Toledo ME, Liu X, Zhao L-R. Stem Cell Factor in Combination With Granulocyte Colony-Stimulating Factor Protects the Brain From Capillary Thrombosis-Induced Ischemic Neuron Loss in a Mouse Model of CADASIL. Front Cell Dev Biol. 2020;8:627733.

63. Schneider A, Krüger C, Steigleder T, Weber D, Pitzer C, Laage R, et al. The hematopoietic factor G-CSF is a neuronal ligand that counteracts programmed cell death and drives neurogenesis. J Clin Invest. 2005;115:2083–98.

64. Schneider A, Kuhn H-G, Schäbitz W-R. A role for G-CSF (granulocyte-colony stimulating factor) in the central nervous system. Cell Cycle Georget Tex. 2005;4:1753–7.

65. Su Y, Cui L, Piao C, Li B, Zhao L-R. The effects of hematopoietic growth factors on neurite outgrowth. PloS One. 2013;8:e75562.

66. Tsai K-J, Tsai Y-C, Shen C-KJ. G-CSF rescues the memory impairment of animal models of Alzheimer’s disease. J Exp Med. 2007;204:1273–80.

67. Barber RC, Edwards MI, Xiao G, Huebinger RM, Diaz-Arrastia R, Wilhelmsen KC, et al. Serum Granulocyte Colony-Stimulating Factor and Alzheimer’s Disease. Dement Geriatr Cogn Disord EXTRA. 2012;2:353–60.

68. Laske C, Sopova K, Hoffmann N, Stransky E, Hagen K, Fallgatter AJ, et al. Stem cell factor plasma levels are decreased in Alzheimer’s disease patients with fast cognitive decline after one-year follow-up period: the Pythia-study. J Alzheimers Dis JAD. 2011;26:39–45.

69. Laske C, Stellos K, Stransky E, Seizer P, Akcay O, Eschweiler GW, et al. Decreased plasma and cerebrospinal fluid levels of stem cell factor in patients with early Alzheimer’s disease. J Alzheimers Dis JAD. 2008;15:451–60.

70. Laske C, Stellos K, Stransky E, Leyhe T, Gawaz M. Decreased plasma levels of granulocyte-colony stimulating factor (G-CSF) in patients with early Alzheimer’s disease. J Alzheimers Dis JAD. 2009;17:115–23.

71. Li B, Gonzalez-Toledo ME, Piao C-S, Gu A, Kelley RE, Zhao L-R. Stem cell factor and granulocyte colony-stimulating factor reduce β-amyloid deposits in the brains of APP/PS1 transgenic mice. Alzheimers Res Ther. 2011;3:8.

72. Keren-Shaul H, Spinrad A, Weiner A, Matcovitch-Natan O, Dvir-Szternfeld R, Ulland TK, et al. A Unique Microglia Type Associated with Restricting Development of Alzheimer’s Disease. Cell. 2017;169:1276–1290.e17.

73. Lee S-H, Meilandt WJ, Xie L, Gandham VD, Ngu H, Barck KH, et al. Trem2 restrains the enhancement of tau accumulation and neurodegeneration by β-amyloid pathology. Neuron. 2021;109:1283–1301.e6.

74. Jankowsky JL, Fadale DJ, Anderson J, Xu GM, Gonzales V, Jenkins NA, et al. Mutant presenilins specifically elevate the levels of the 42 residue beta-amyloid peptide in vivo: evidence for augmentation of a 42-specific gamma secretase. Hum Mol Genet. 2004;13:159–70.

75. Benjamini Y, Hochberg Y. Controlling the False Discovery Rate: A Practical and Powerful Approach to Multiple Testing. J R Stat Soc Ser B Methodol. 1995;57:289–300.

76. Goedhart J, Luijsterburg MS. VolcaNoseR is a web app for creating, exploring, labeling and sharing volcano plots. Sci Rep. 2020;10:20560.

77. Doncheva NT, Morris JH, Gorodkin J, Jensen LJ. Cytoscape StringApp: Network Analysis and Visualization of Proteomics Data. J Proteome Res. 2019;18:623–32.

78. Shannon P, Markiel A, Ozier O, Baliga NS, Wang JT, Ramage D, et al. Cytoscape: a software environment for integrated models of biomolecular interaction networks. Genome Res. 2003;13:2498–504.

79. Haage V, Semtner M, Vidal RO, Hernandez DP, Pong WW, Chen Z, et al. Comprehensive gene expression meta-analysis identifies signature genes that distinguish microglia from peripheral monocytes/macrophages in health and glioma. Acta Neuropathol Commun. 2019;7:20.

80. Hohsfield LA, Tsourmas KI, Ghorbanian Y, Syage AR, Jin Kim S, Cheng Y, et al. MAC2 is a long-lasting marker of peripheral cell infiltrates into the mouse CNS after bone marrow transplantation and coronavirus infection. Glia. 2022;70:875–91.

81. Sevenich L. Brain-Resident Microglia and Blood-Borne Macrophages Orchestrate Central Nervous System Inflammation in Neurodegenerative Disorders and Brain Cancer. Front Immunol [Internet]. 2018 [cited 2024 Jun 2];9. Available from: https://www.frontiersin.org/journals/immunology/articles/10.3389/fimmu.2018.00697/full

82. Xia C, Braunstein Z, Toomey AC, Zhong J, Rao X. S100 Proteins As an Important Regulator of Macrophage Inflammation. Front Immunol [Internet]. 2018 [cited 2022 Sep 8];8. Available from: https://www.frontiersin.org/articles/10.3389/fimmu.2017.01908

83. Gray E, Thomas TL, Betmouni S, Scolding N, Love S. Elevated activity and microglial expression of myeloperoxidase in demyelinated cerebral cortex in multiple sclerosis. Brain Pathol Zurich Switz. 2008;18:86–95.

84. Odink K, Cerletti N, Brüggen J, Clerc RG, Tarcsay L, Zwadlo G, et al. Two calcium-binding proteins in infiltrate macrophages of rheumatoid arthritis. Nature. 1987;330:80–2.

85. Hickman SE, Kingery ND, Ohsumi T, Borowsky M, Wang L, Means TK, et al. The Microglial Sensome Revealed by Direct RNA Sequencing. Nat Neurosci. 2013;16:1896–905.

86. McFarland HI, Nahill SR, Maciaszek JW, Welsh RM. CD11b (Mac-1): a marker for CD8+ cytotoxic T cell activation and memory in virus infection. J Immunol Baltim Md 1950. 1992;149:1326–33.

87. Butovsky O, Weiner HL. Microglial signatures and their role in health and disease. Nat Rev Neurosci. 2018;19:622–35.

88. Deczkowska A, Keren-Shaul H, Weiner A, Colonna M, Schwartz M, Amit I. Disease-Associated Microglia: A Universal Immune Sensor of Neurodegeneration. Cell. 2018;173:1073–81.

89. Rangaraju S, Dammer EB, Raza SA, Rathakrishnan P, Xiao H, Gao T, et al. Identification and therapeutic modulation of a pro-inflammatory subset of disease-associated-microglia in Alzheimer’s disease. Mol Neurodegener. 2018;13:24.

90. Zhang GX, Li J, Ventura E, Rostami A. Parenchymal microglia of naïve adult C57BL/6J mice express high levels of B7.1, B7.2, and MHC class II. Exp Mol Pathol. 2002;73:35–45.

91. Ford AL, Goodsall AL, Hickey WF, Sedgwick JD. Normal adult ramified microglia separated from other central nervous system macrophages by flow cytometric sorting. Phenotypic differences defined and direct ex vivo antigen presentation to myelin basic protein-reactive CD4+ T cells compared. J Immunol Baltim Md 1950. 1995;154:4309–21.

92. Cristóvão JS, Gomes CM. S100 Proteins in Alzheimer’s Disease. Front Neurosci. 2019;13:463.

93. Kummer MP, Vogl T, Axt D, Griep A, Vieira-Saecker A, Jessen F, et al. Mrp14 deficiency ameliorates amyloid β burden by increasing microglial phagocytosis and modulation of amyloid precursor protein processing. J Neurosci Off J Soc Neurosci. 2012;32:17824–9.

94. Grabert K, Michoel T, Karavolos MH, Clohisey S, Baillie JK, Stevens MP, et al. Microglial brain region-dependent diversity and selective regional sensitivities to aging. Nat Neurosci. 2016;19:504–16.

95. Chen Y, Colonna M. Microglia in Alzheimer’s disease at single-cell level. Are there common patterns in humans and mice? J Exp Med. 2021;218:e20202717.

96. Hammond TR, Dufort C, Dissing-Olesen L, Giera S, Young A, Wysoker A, et al. Single-Cell RNA Sequencing of Microglia throughout the Mouse Lifespan and in the Injured Brain Reveals Complex Cell-State Changes. Immunity. 2019;50:253–271.e6.

97. Orre M, Kamphuis W, Osborn LM, Melief J, Kooijman L, Huitinga I, et al. Acute isolation and transcriptome characterization of cortical astrocytes and microglia from young and aged mice. Neurobiol Aging. 2014;35:1– 14.

98. Wang H. Microglia Heterogeneity in Alzheimer’s Disease: Insights From Single-Cell Technologies. Front Synaptic Neurosci [Internet]. 2021 [cited 2022 Dec 17];13. Available from: https://www.frontiersin.org/articles/10.3389/fnsyn.2021.773590

99. Kenkhuis B, Somarakis A, Kleindouwel LRT, van Roon-Mom WMC, Höllt T, van der Weerd L. Co-expression patterns of microglia markers Iba1, TMEM119 and P2RY12 in Alzheimer’s disease. Neurobiol Dis. 2022;167:105684.

100. Boyd TD, Bennett SP, Mori T, Governatori N, Runfeldt M, Norden M, et al. GM-CSF upregulated in rheumatoid arthritis reverses cognitive impairment and amyloidosis in Alzheimer mice. J Alzheimers Dis JAD. 2010;21:507–18.

101. Kiyota T, Machhi J, Lu Y, Dyavarshetty B, Nemati M, Yokoyama I, et al. Granulocyte-macrophage colony-stimulating factor neuroprotective activities in Alzheimer’s disease mice. J Neuroimmunol. 2018;319:80–92.

102. Rangaraju S, Raza SA, Li NX, Betarbet R, Dammer EB, Duong D, et al. Differential Phagocytic Properties of CD45low Microglia and CD45high Brain Mononuclear Phagocytes-Activation and Age-Related Effects. Front Immunol. 2018;9:405.

103. Hohsfield LA, Humpel C. Intravenous Infusion of Monocytes Isolated from 2-Week-Old Mice Enhances Clearance of Beta-Amyloid Plaques in an Alzheimer Mouse Model. PLOS ONE. 2015;10:e0121930.

104. Zuroff L, Daley D, Black KL, Koronyo-Hamaoui M. Clearance of cerebral Aβ in Alzheimer’s disease: reassessing the role of microglia and monocytes. Cell Mol Life Sci CMLS. 2017;74:2167–201.

105. Vogel DYS, Kooij G, Heijnen PDAM, Breur M, Peferoen LAN, van der Valk P, et al. GM-CSF promotes migration of human monocytes across the blood brain barrier. Eur J Immunol. 2015;45:1808–19.

106. Potter H, Woodcock JH, Boyd TD, Coughlan CM, O’Shaughnessy JR, Borges MT, et al. Safety and efficacy of sargramostim (GM-CSF) in the treatment of Alzheimer’s disease. Alzheimers Dement Transl Res Clin Interv. 2021;7:e12158.

107. Wang S, Song R, Wang Z, Jing Z, Wang S, Ma J. S100A8/A9 in Inflammation. Front Immunol [Internet]. 2018 [cited 2022 Dec 8];9. Available from: https://www.frontiersin.org/articles/10.3389/fimmu.2018.01298

108. Gong Y, Hart E, Shchurin A, Hoover-Plow J. Inflammatory macrophage migration requires MMP-9 activation by plasminogen in mice. J Clin Invest. 2008;118:3012–24.

109. Gross SR, Sin CGT, Barraclough R, Rudland PS. Joining S100 proteins and migration: for better or for worse, in sickness and in health. Cell Mol Life Sci CMLS. 2014;71:1551–79.

110. Jang E, Lee S, Kim J-H, Kim J-H, Seo J-W, Lee W-H, et al. Secreted protein lipocalin-2 promotes microglial M1 polarization. FASEB J Off Publ Fed Am Soc Exp Biol. 2013;27:1176–90.

111. Kang H, Shin HJ, An HS, Jin Z, Lee JY, Lee J, et al. Role of Lipocalin-2 in Amyloid-Beta Oligomer-Induced Mouse Model of Alzheimer’s Disease. Antioxid Basel Switz. 2021;10:1657.

112. Lee S, Kim J-H, Kim J-H, Seo J-W, Han H-S, Lee W-H, et al. Lipocalin-2 Is a chemokine inducer in the central nervous system: role of chemokine ligand 10 (CXCL10) in lipocalin-2-induced cell migration. J Biol Chem. 2011;286:43855–70.

113. Ehrchen JM, Sunderkötter C, Foell D, Vogl T, Roth J. The endogenous Toll–like receptor 4 agonist S100A8/S100A9 (calprotectin) as innate amplifier of infection, autoimmunity, and cancer. J Leukoc Biol. 2009;86:557–66.

114. He Z, Riva M, Björk P, Swärd K, Mörgelin M, Leanderson T, et al. CD14 Is a Co-Receptor for TLR4 in the S100A9-Induced Pro-Inflammatory Response in Monocytes. PLoS ONE. 2016;11:e0156377.

115. Vogl T, Ludwig S, Goebeler M, Strey A, Thorey IS, Reichelt R, et al. MRP8 and MRP14 control microtubule reorganization during transendothelial migration of phagocytes. Blood. 2004;104:4260–8.

116. Chang K-A, Kim HJ, Suh Y-H. The role of S100a9 in the pathogenesis of Alzheimer’s disease: the therapeutic effects of S100a9 knockdown or knockout. Neurodegener Dis. 2012;10:27–9.

117. Pansieri J, Iashchishyn IA, Fakhouri H, Ostojić L, Malisauskas M, Musteikyte G, et al. Templating S100A9 amyloids on Aβ fibrillar surfaces revealed by charge detection mass spectrometry, microscopy, kinetic and microfluidic analyses. Chem Sci. 2020;11:7031–9.

118. Wang C, Klechikov AG, Gharibyan AL, Wärmländer SKTS, Jarvet J, Zhao L, et al. The role of pro-inflammatory S100A9 in Alzheimer’s disease amyloid-neuroinflammatory cascade. Acta Neuropathol (Berl). 2014;127:507–22.

119. Russo A, Schürmann H, Brandt M, Scholz K, Matos ALL, Grill D, et al. Alarming and Calming: Opposing Roles of S100A8/S100A9 Dimers and Tetramers on Monocytes. Adv Sci Weinh Baden-Wurtt Ger. 2022;e2201505.

120. Karlstetter M, Walczak Y, Weigelt K, Ebert S, Van den Brulle J, Schwer H, et al. The novel activated microglia/macrophage WAP domain protein, AMWAP, acts as a counter-regulator of proinflammatory response. J Immunol Baltim Md 1950. 2010;185:3379–90.

121. Camilli C, Hoeh AE, De Rossi G, Moss SE, Greenwood J. LRG1: an emerging player in disease pathogenesis. J Biomed Sci. 2022;29:6.

122. Ikeda N, Asano K, Kikuchi K, Uchida Y, Ikegami H, Takagi R, et al. Emergence of immunoregulatory Ym1+Ly6Chi monocytes during recovery phase of tissue injury. Sci Immunol. 2018;3:eaat0207.

123. Kang Q, Li L, Pang Y, Zhu W, Meng L. An update on Ym1 and its immunoregulatory role in diseases. Front Immunol [Internet]. 2022 [cited 2023 Jan 1];13. Available from: https://www.frontiersin.org/articles/10.3389/fimmu.2022.891220

124. Liu K, Tian L-X, Tang X, Wang J, Tang W-Q, Ma Z-F, et al. Neutrophilic granule protein (NGP) attenuates lipopolysaccharide-induced inflammatory responses and enhances phagocytosis of bacteria by macrophages. Cytokine. 2020;128:155001.

125. Makita N, Hizukuri Y, Yamashiro K, Murakawa M, Hayashi Y. IL-10 enhances the phenotype of M2 macrophages induced by IL-4 and confers the ability to increase eosinophil migration. Int Immunol. 2015;27:131–41.

126. Liu N, Jiang Y, Chung JY, Li Y, Yu Z, Kim JW, et al. Annexin A2 Deficiency Exacerbates Neuroinflammation and Long-Term Neurological Deficits after Traumatic Brain Injury in Mice. Int J Mol Sci. 2019;20:6125.

127. Morimoto M, Nakano T, Egashira S, Irie K, Matsuyama K, Wada M, et al. Haptoglobin Regulates Macrophage/Microglia-Induced Inflammation and Prevents Ischemic Brain Damage Via Binding to HMGB1. J Am Heart Assoc. 2022;11:e024424.

128. McArthur S, Cristante E, Paterno M, Christian H, Roncaroli F, Gillies GE, et al. Annexin A1: a central player in the anti-inflammatory and neuroprotective role of microglia. J Immunol Baltim Md 1950. 2010;185:6317– 28.

129. Aslanidis A, Karlstetter M, Scholz R, Fauser S, Neumann H, Fried C, et al. Activated microglia/macrophage whey acidic protein (AMWAP) inhibits NFκB signaling and induces a neuroprotective phenotype in microglia. J Neuroinflammation. 2015;12:77.

130. Ries M, Loiola R, Shah UN, Gentleman SM, Solito E, Sastre M. The anti-inflammatory Annexin A1 induces the clearance and degradation of the amyloid-β peptide. J Neuroinflammation. 2016;13:234.

131. Ries M, Watts H, Mota BC, Lopez MY, Donat CK, Baxan N, et al. Annexin A1 restores cerebrovascular integrity concomitant with reduced amyloid-β and tau pathology. Brain J Neurol. 2021;144:1526–41.

132. Xia Q, Li X, Zhou H, Zheng L, Shi J. S100A11 protects against neuronal cell apoptosis induced by cerebral ischemia via inhibiting the nuclear translocation of annexin A1. Cell Death Dis. 2018;9:657.

133. Tian Z-Y, Wang C-Y, Wang T, Li Y-C, Wang Z-Y. Glial S100A6 Degrades β-amyloid Aggregation through Targeting Competition with Zinc Ions. Aging Dis. 2019;10:756–69.

134. Garland P, Morton MJ, Haskins W, Zolnourian A, Durnford A, Gaastra B, et al. Haemoglobin causes neuronal damage in vivo which is preventable by haptoglobin. Brain Commun. 2020;2:fcz053.

135. Graff-Radford J, Lesnick T, Rabinstein AA, Gunter J, Aakre J, Przybelski SA, et al. Cerebral microbleed incidence, relationship to amyloid burden: The Mayo Clinic Study of Aging. Neurology. 2020;94:e190–9.

136. Yerbury JJ, Kumita JR, Meehan S, Dobson CM, Wilson MR. α2-Macroglobulin and Haptoglobin Suppress Amyloid Formation by Interacting with Prefibrillar Protein Species *. J Biol Chem. 2009;284:4246–54.

137. Paudel YN, Angelopoulou E, Piperi C, Othman I, Aamir K, Shaikh MF. Impact of HMGB1, RAGE, and TLR4 in Alzheimer’s Disease (AD): From Risk Factors to Therapeutic Targeting. Cells. 2020;9:383.

138. Takata K, Kitamura Y, Tsuchiya D, Kawasaki T, Taniguchi T, Shimohama S. High mobility group box protein-1 inhibits microglial Abeta clearance and enhances Abeta neurotoxicity. J Neurosci Res. 2004;78:880–91.

139. Vafadari B, Salamian A, Kaczmarek L. MMP-9 in translation: from molecule to brain physiology, pathology, and therapy. J Neurochem. 2016;139:91–114.

140. Miners JS, Baig S, Palmer J, Palmer LE, Kehoe PG, Love S. Abeta-degrading enzymes in Alzheimer’s disease. Brain Pathol Zurich Switz. 2008;18:240–52.

141. Yan P, Hu X, Song H, Yin K, Bateman RJ, Cirrito JR, et al. Matrix metalloproteinase-9 degrades amyloid-beta fibrils in vitro and compact plaques in situ. J Biol Chem. 2006;281:24566–74.

142. Fragkouli A, Tsilibary EC, Tzinia AK. Neuroprotective role of MMP-9 overexpression in the brain of Alzheimer’s 5xFAD mice. Neurobiol Dis. 2014;70:179–89.

143. Dar NJ, Glazner GW. Deciphering the neuroprotective and neurogenic potential of soluble amyloid precursor protein alpha (sAPPα). Cell Mol Life Sci CMLS. 2020;77:2315–30.

144. Mockett BG, Richter M, Abraham WC, Müller UC. Therapeutic Potential of Secreted Amyloid Precursor Protein APPsα. Front Mol Neurosci. 2017;10:30.

145. Das N, Raymick J, Sarkar S. Role of metals in Alzheimer’s disease. Metab Brain Dis. 2021;36:1627–39.

146. Wang L, Yin Y-L, Liu X-Z, Shen P, Zheng Y-G, Lan X-R, et al. Current understanding of metal ions in the pathogenesis of Alzheimer’s disease. Transl Neurodegener. 2020;9:10.

147. Hunter M, Spiller KJ, Dominique MA, Xu H, Hunter FW, Fang TC, et al. Microglial transcriptome analysis in the rNLS8 mouse model of TDP-43 proteinopathy reveals discrete expression profiles associated with neurodegenerative progression and recovery. Acta Neuropathol Commun. 2021;9:140.

148. Smyth L, Murray H, Hill M, Leeuwen E, Highet B, Magon N, et al. Neutrophil-vascular interactions drive myeloperoxidase accumulation in the brain in Alzheimer’s disease. Acta Neuropathol Commun. 2022;10.

149. Zenaro E, Pietronigro E, Bianca VD, Piacentino G, Marongiu L, Budui S, et al. Neutrophils promote Alzheimer’s disease–like pathology and cognitive decline via LFA-1 integrin. Nat Med. 2015;21:880–6.

150. Meyer-Luehmann M, Prinz M. Myeloid cells in Alzheimer’s disease: culprits, victims or innocent bystanders? Trends Neurosci. 2015;38:659–68.

151. Gordon B, Blazey T, Benzinger T. Regional variability in Alzheimer’s disease biomarkers. Future Neurol. 2014;9:131–4.

