## Supplementary figures and images for "Single cell RNA sequencing reveals immunomodulatory effects of stem cell factor and granulocyte colony-stimulating factor treatment in the brains of aged APP/PS1 mice"

### Supplemental Fig 1

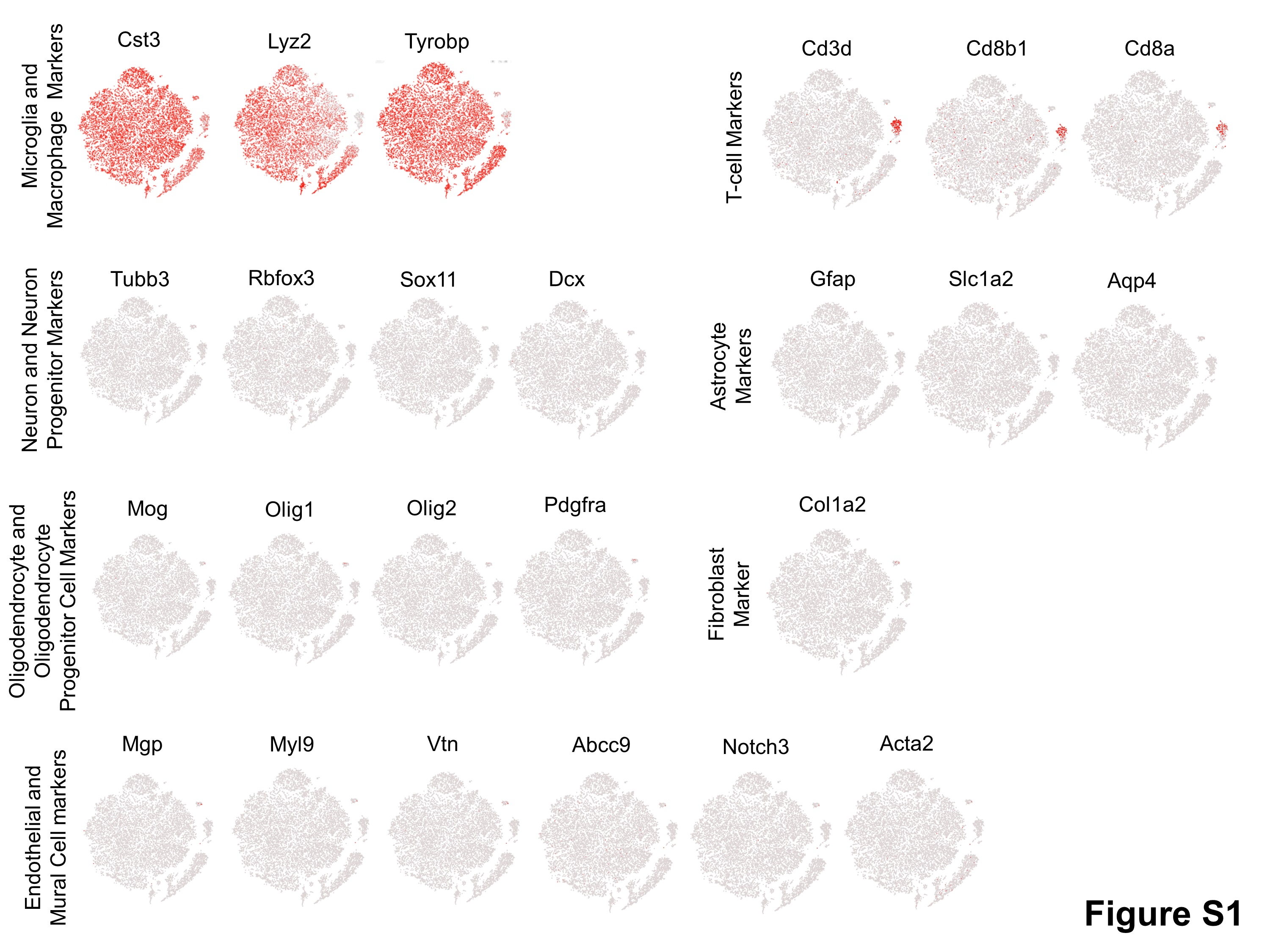

### Supplemental Fig 2

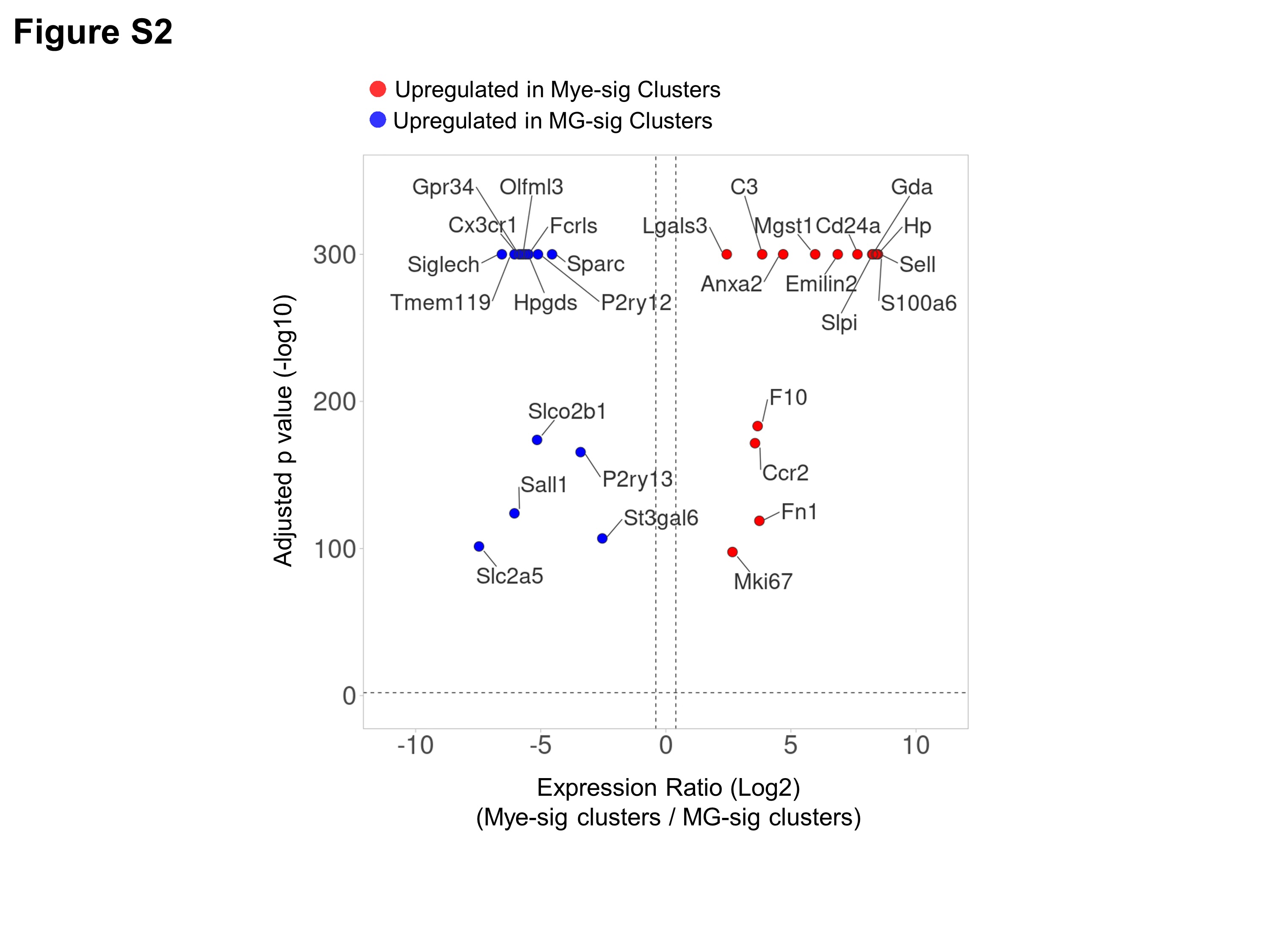

### Supplemental Fig 3

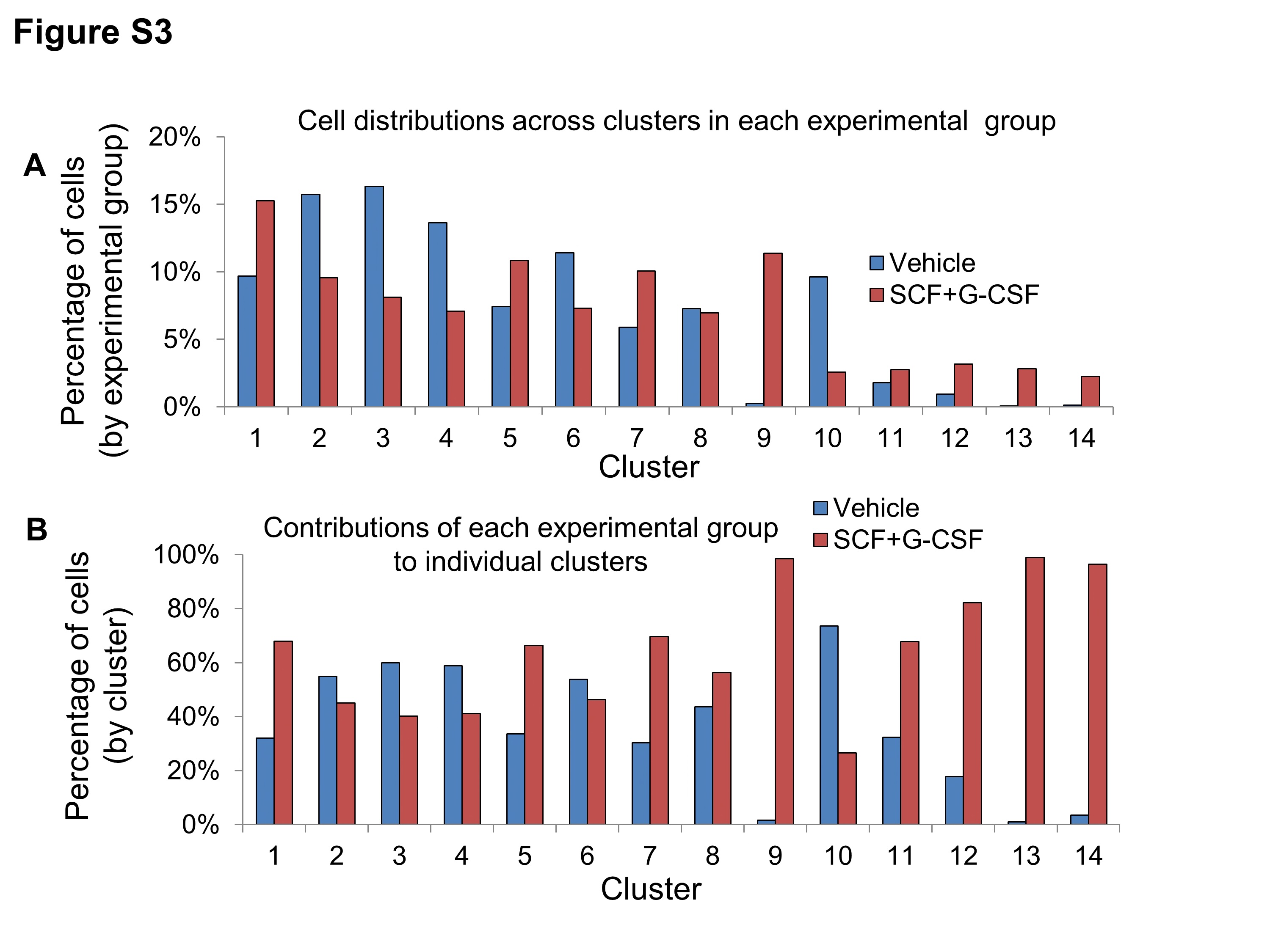

### Supplemental Fig 4

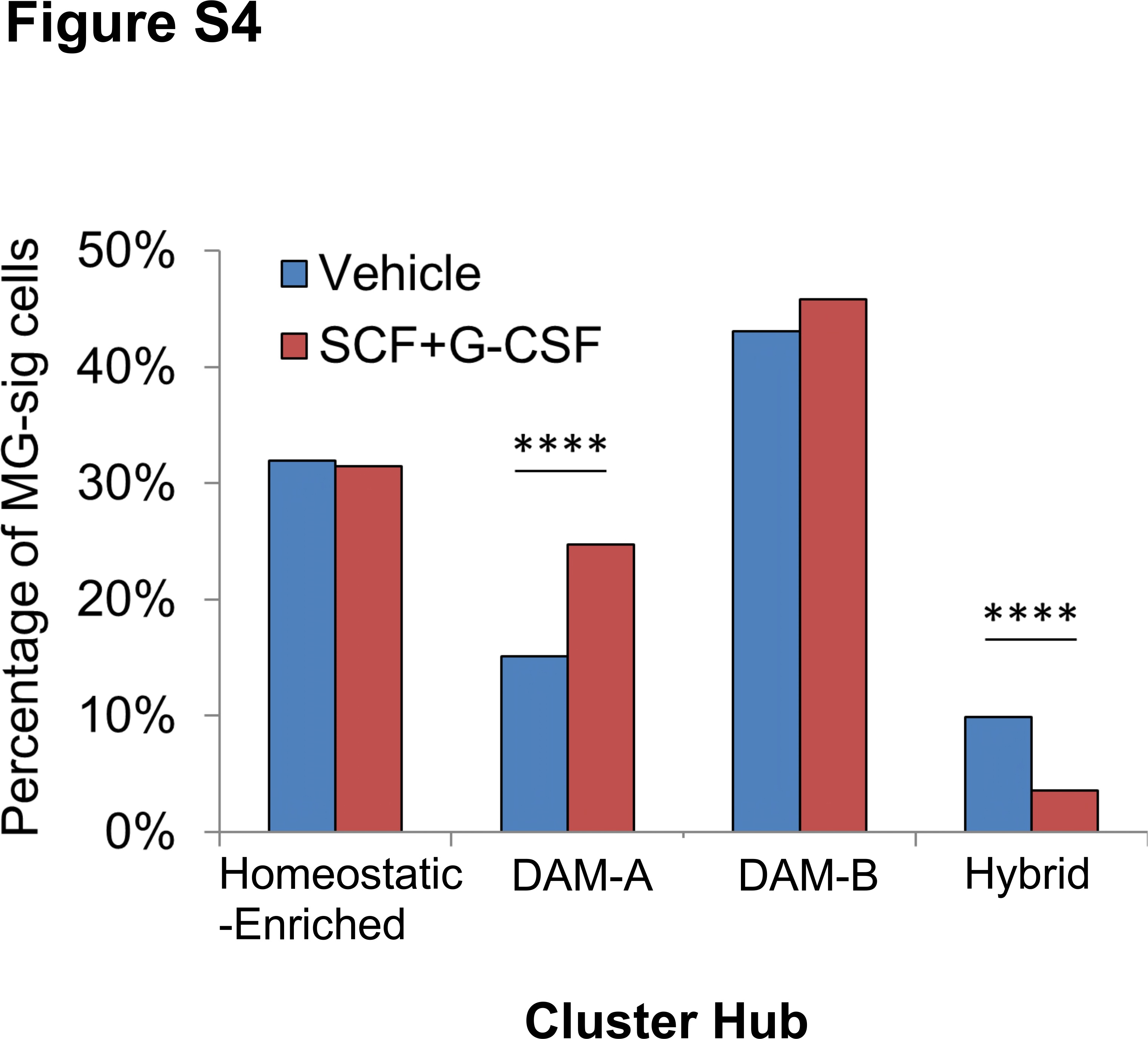

### Supplemental Fig 5

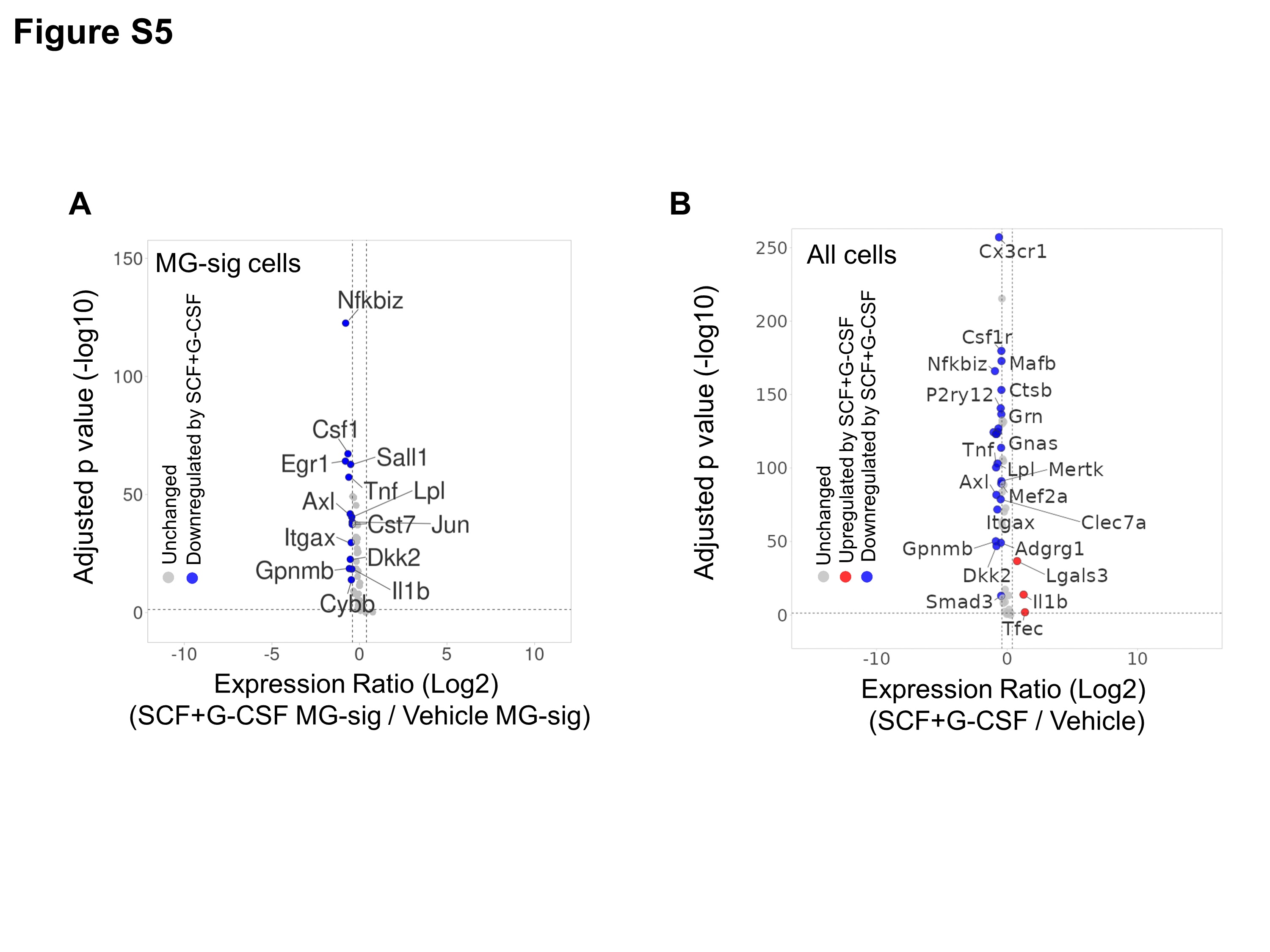

### Supplemental Fig 6

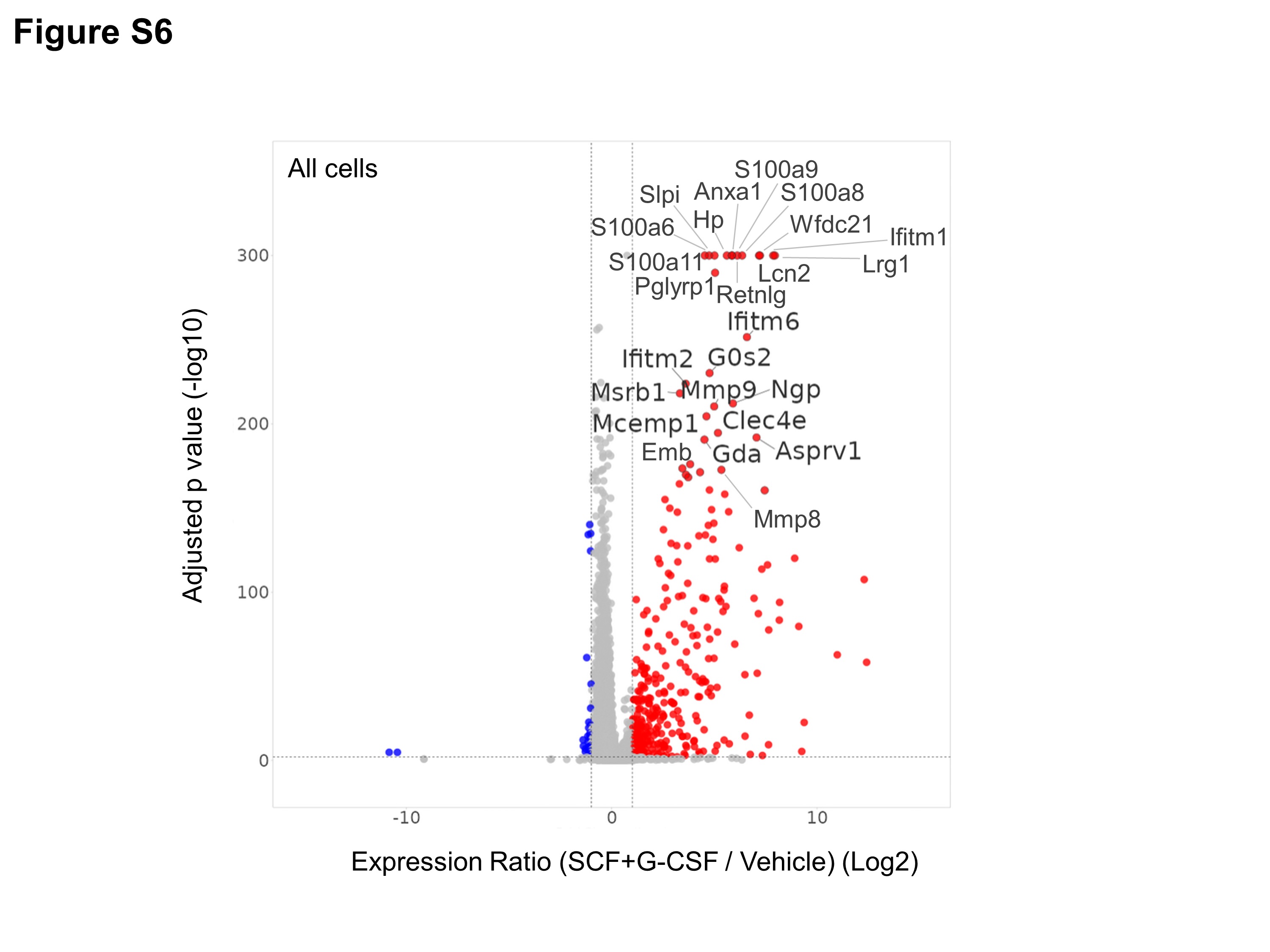

### Supplemental Fig 7

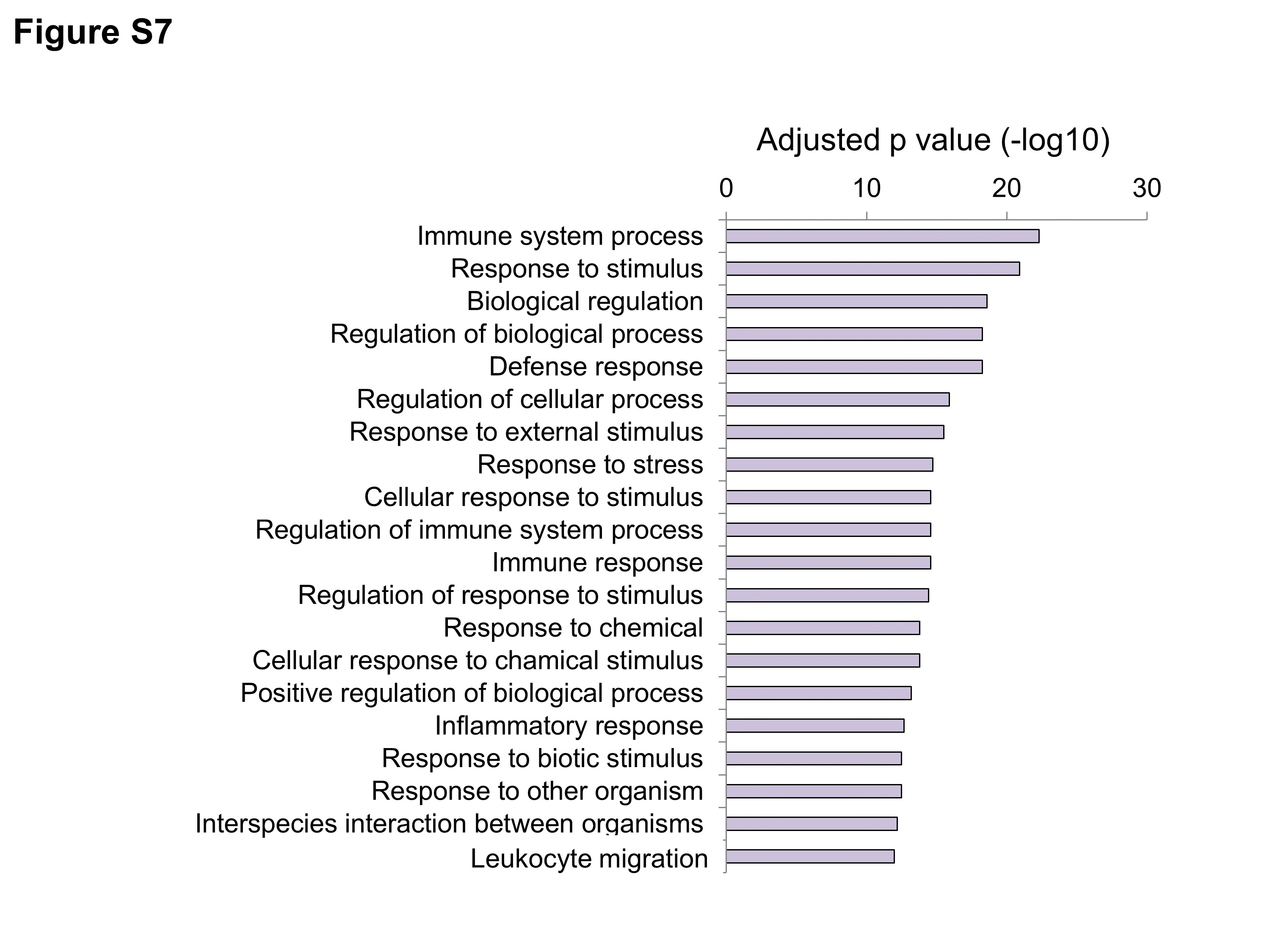

### Supplemental Fig 8

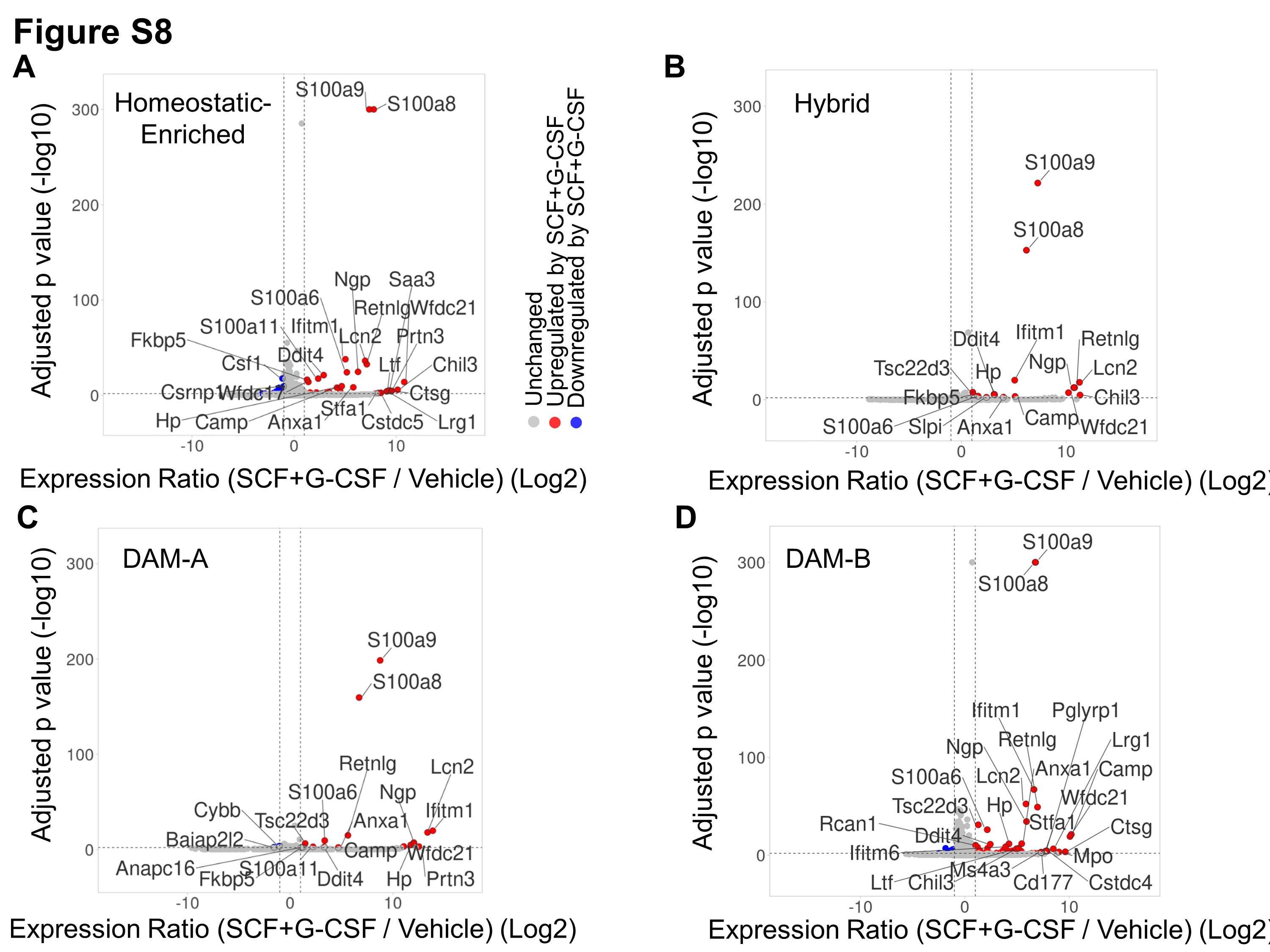

### Supplemental Fig 9

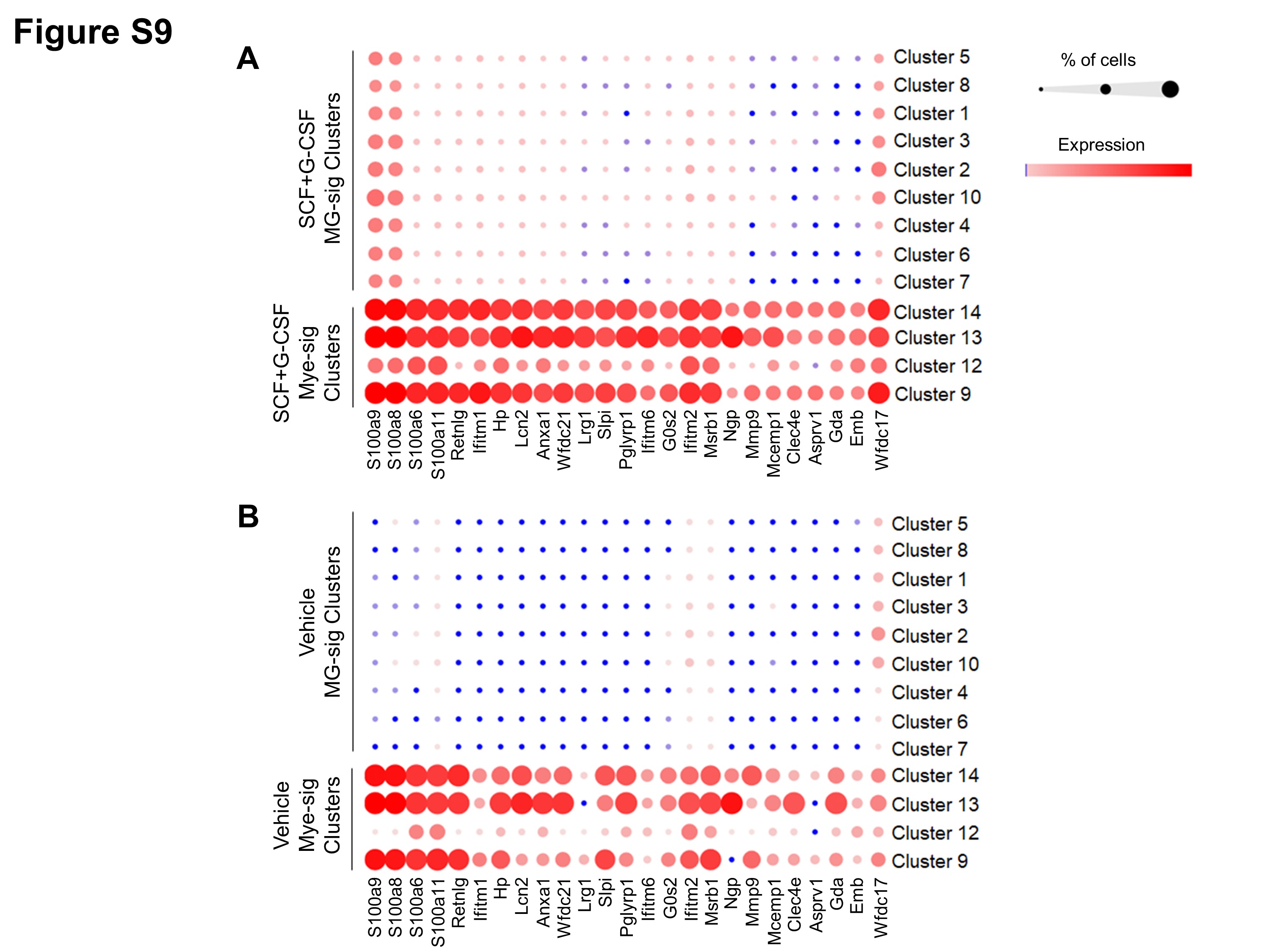

### Supplemental Fig 10

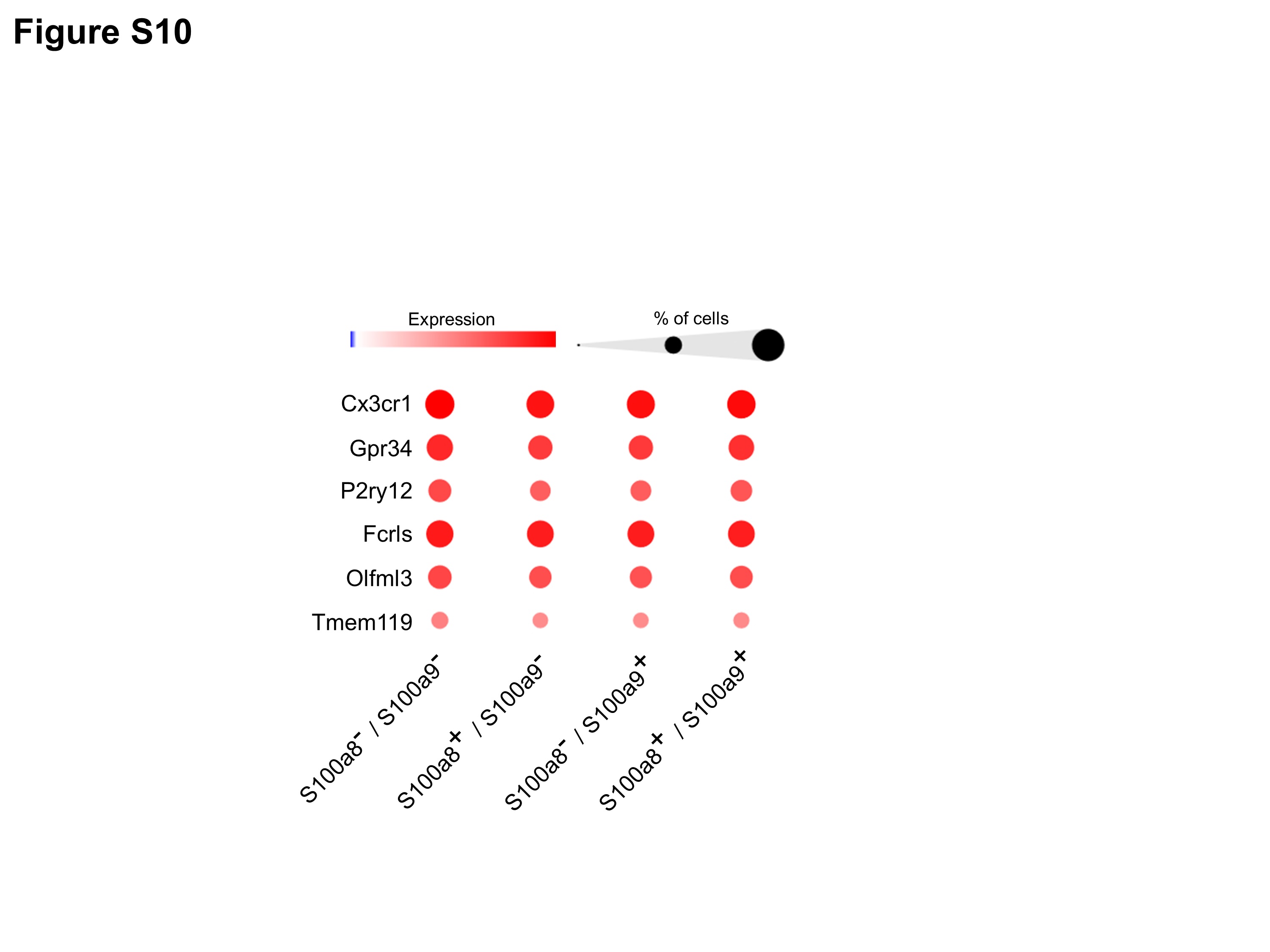

### Supplemental Fig 11

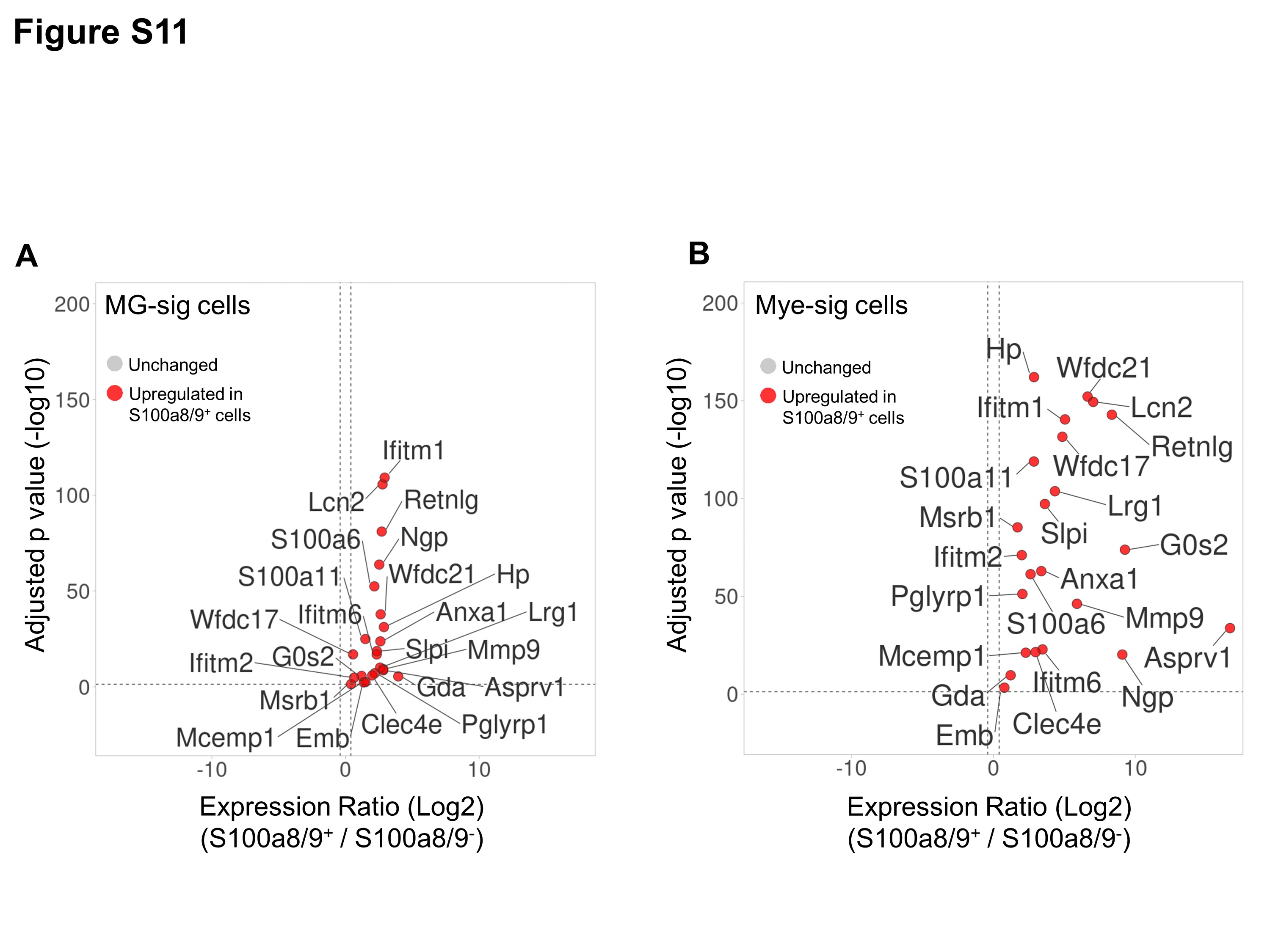

### Supplemental Fig 12

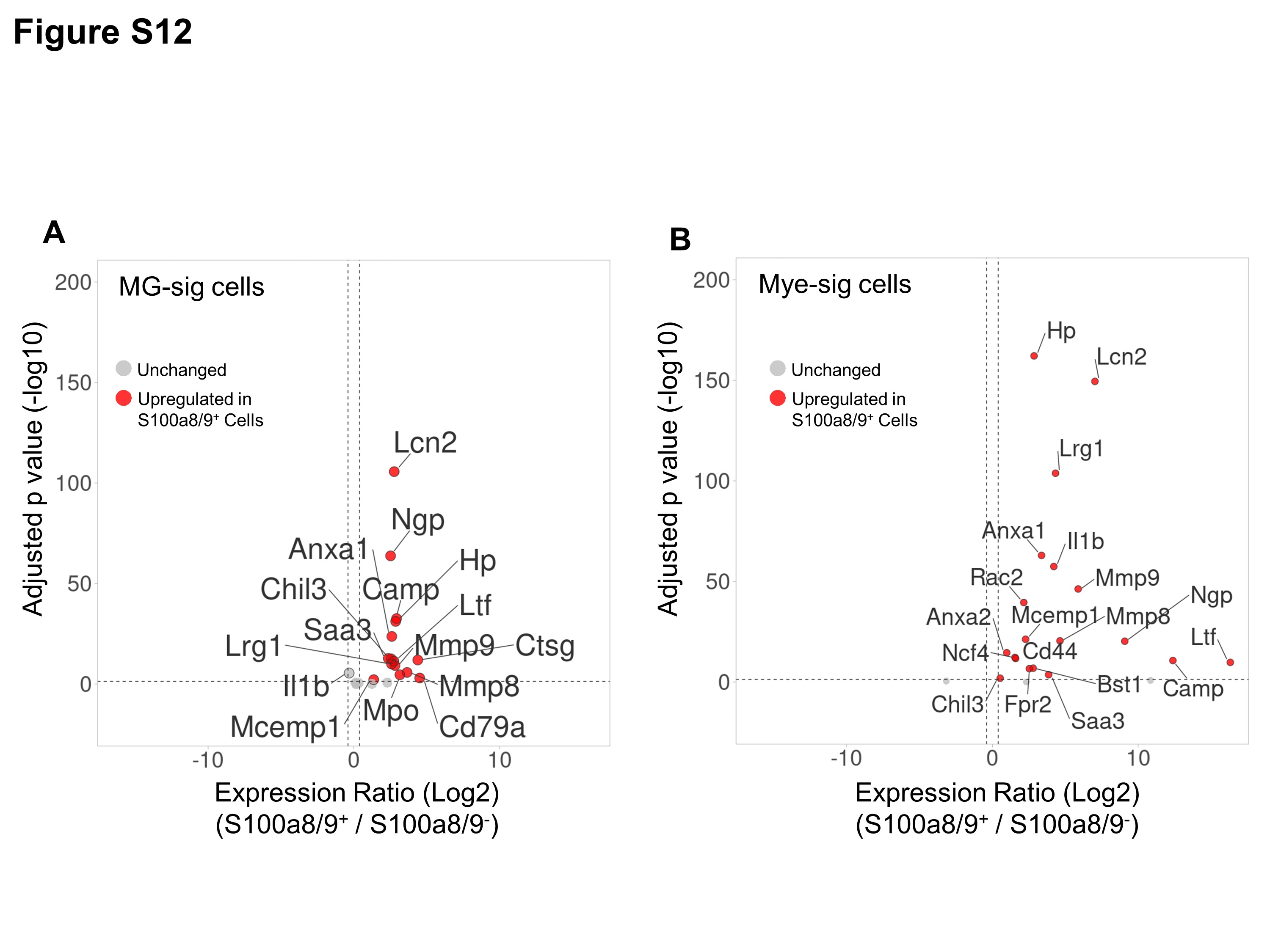

### Supplemental Fig 13

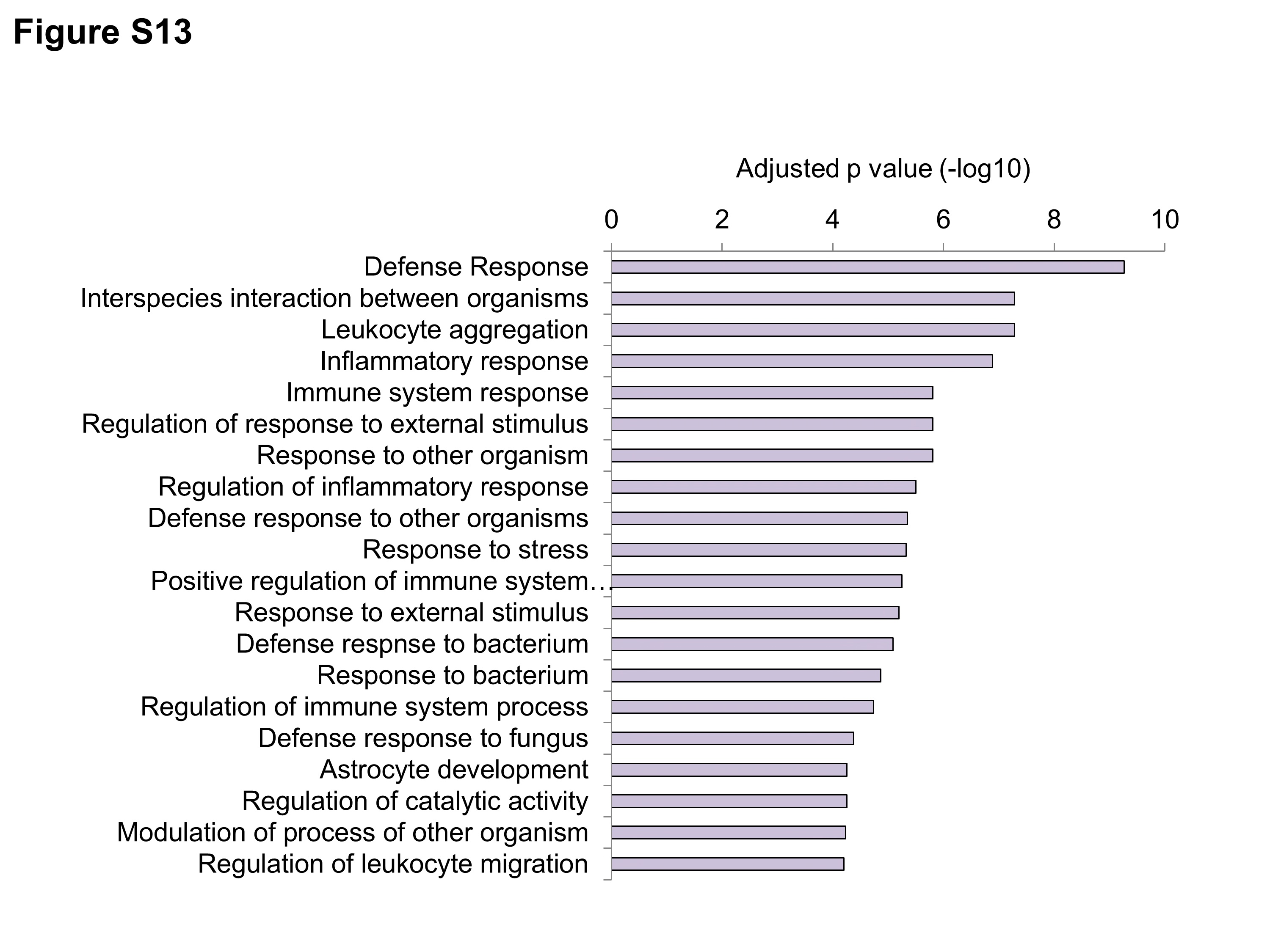

### Supplemental Table 1

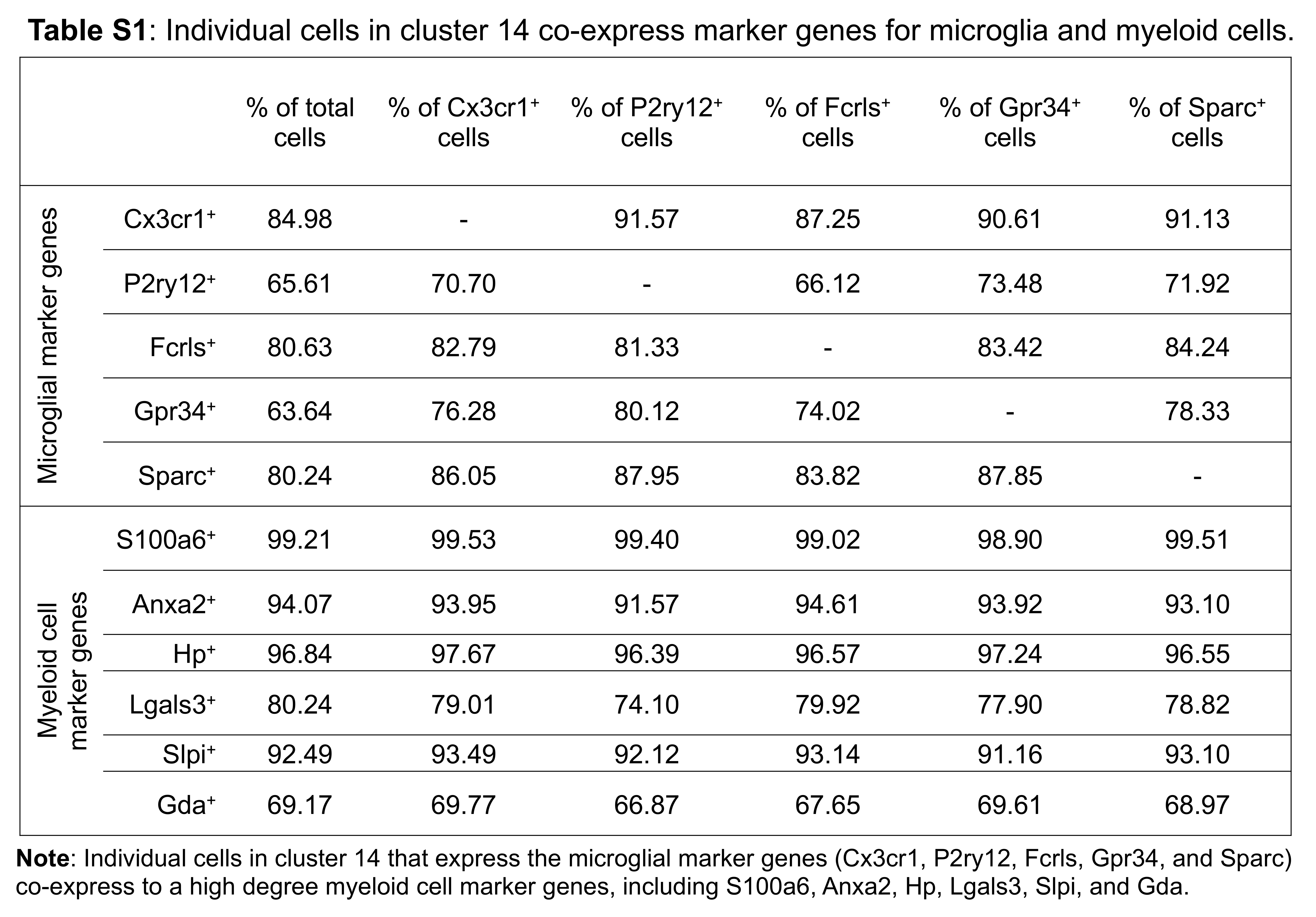

### Supplemental Table 2

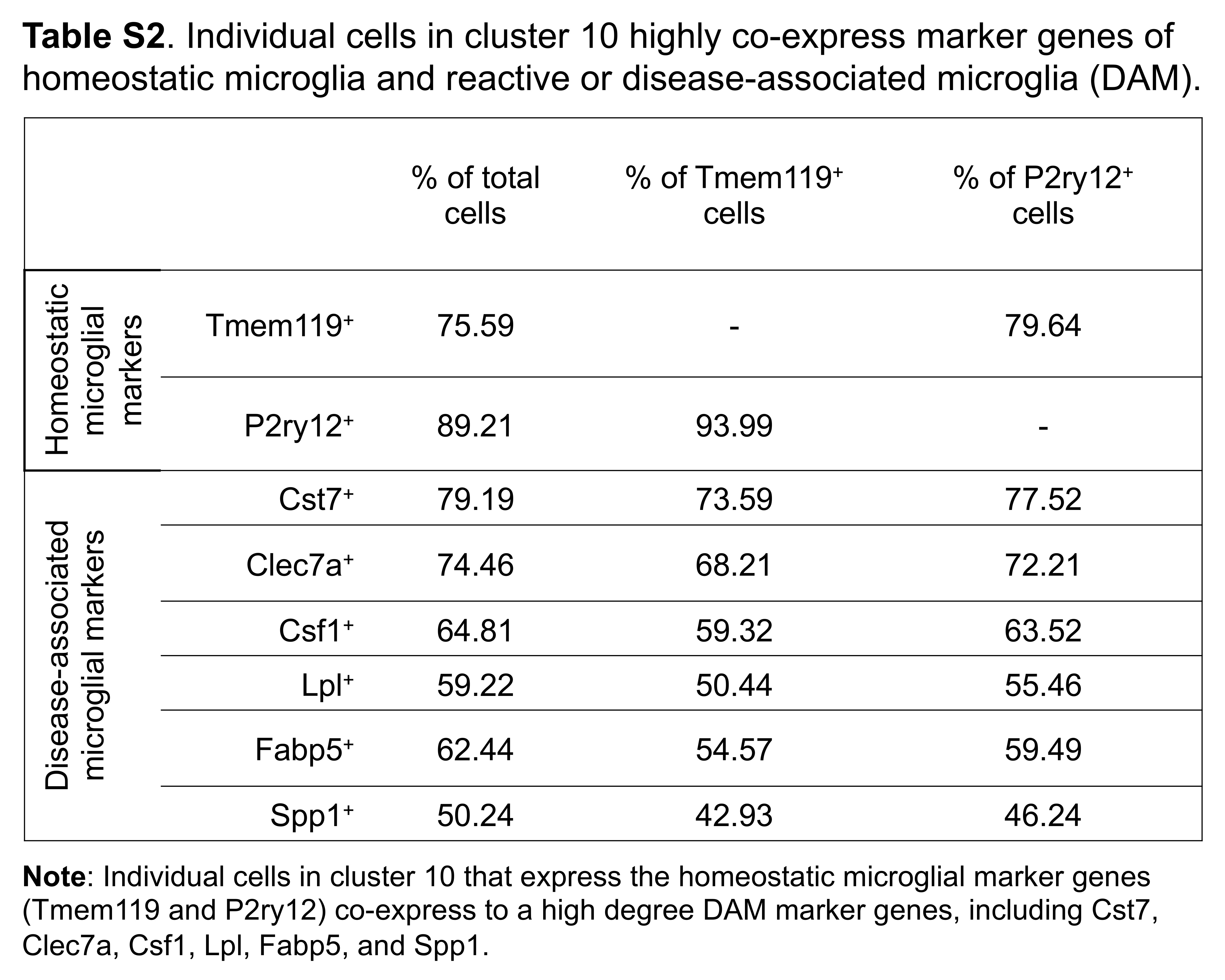
