## Supplemental Figure legends for "Single cell RNA sequencing reveals immunomodulatory effects of stem cell factor and granulocyte colony-stimulating factor treatment in the brains of aged APP/PS1 mice"

**Figure S1. Cell clusters highly express marker genes of microglia and macrophages but not marker genes of off-target cells in the brain.** Expression (in red) of marker genes of various cell classes across tSNE plots indicate that clusters 1-10 and 12-14 selectively express microglial cell and myeloid cell marker genes (Cst3, Lyz2, Tyropb), while cluster 11 selectively expresses T-cell marker genes. None of the clusters prominently expresses marker genes of other primary cell classes in the brain, including neurons, neuron progenitor cells, astrocytes, oligodendrocytes, oligodendrocyte progenitor cells, endothelial cells, mural cells, and fibroblasts.

**Figure S2. Myeloid cell marker genes and microglial marker genes are respectively enriched in Mye-sig and MG-sig cell clusters.**  A volcano plot of microglial and myeloid cell marker genes highlights DEGs in MG-sig clusters relative to Mye-sig clusters (clusters 1-8,10 vs. clusters 9, 12-13, respectively). Genes upregulated in Mye-sig clusters are shown in red. Genes upregulated in MG-sig clusters are shown in blue. False discovery rate-corrected *p* values (-log10) are shown on the y-axis. The log2-transformed expression ratios are computed as expression levels in Mye-sig clusters relative to those in MG-sig clusters and are plotted on the x-axis. Dotted lines indicate significance thresholds. Note, *p* values are restricted to 300 decimal places.

**Figure S3. The percentage of cells contained in individual clusters across experimental groups.** (**A**) The percentage contribution by each cluster to each experimental group. Note, the summed percentage of cells in all clusters for each individual experimental group is 100%. (**B**) The percentage contribution by each experimental group to individual clusters. Note, the summed percentage of cells in both experimental groups combined for each individual cluster is 100%.

**Figure S4. SCF+G-CSF treatment alters the reactive profiles of MG-sig cells.** SCF+G-CSF treatment increases the percentage of MG-sig cells in the DAM-A hub and reduces the percentage of cells in the hybrid hub (cluster 10). **** p < 0.0001.

**Figure S5.** **SCF+G-CSF treatment modestly alters gene sets associated with homeostatic microglia, DAM, and inflammatory microglia .** Volcano plots highlight homeostatic, DAM, and inflammatory gene sets that are differentially expressed in SCF+G-CSF treatment in MG-sig clusters pooled together (1-8, 10 clusters) (**A**), and in all clusters pooled together (**B**). Genes upregulated by SCF+G-CSF treatment are shown in red. Genes downregulated by SCF+G-CSF treatment are shown in blue. False discovery rate-corrected *p* values (-log10) are shown on the y-axis. The log2-transformed expression ratios are computed as expression levels in the SCF+G-CSF-treated mice relative to those in the vehicle control group. The log2-transformed expression ratios are plotted on the x-axis. Dotted lines indicate significance thresholds.

**Figure S6.** **Transcriptome-wide responses to SCF+G-CSF treatment.** A volcano plot highlights the genes differentially expressed by SCF+G-CSF treatment in all clusters pooled together. Genes upregulated by SCF+G-CSF treatment are shown in red. Genes downregulated by SCF+G-CSF treatment are shown in blue. False discovery rate-corrected *p* values (-log10) are shown on the y-axis. The log2-transformed expression ratios are computed as expression levels in the SCF+G-CSF-treated mice relative to those in the vehicle control group. The log2-transformed expression ratios are plotted on the x-axis. Dotted lines indicate significance thresholds. Note, *p* values are restricted to 300 decimal places. The top 25 differentially expressed genes are labeled.

**Figure S7. Gene ontology analysis of all DEGs upregulated by SCF+G-CSF treatment.** Gene ontology analysis of all differentially expressed genes (DEGs) following SCF+G-CSF treatment reveals biological functions altered by the SCF+G-CSF treatment. These functions include regulation of the immune system, responses to stress and inflammation, and leukocyte migration.

**Figure S8. Transcriptome-wide responses to SCF+G-CSF treatment in MG-sig cluster hubs.** (**A**-**D**) Volcano plots highlight the genes differentially expressed in SCF+G-CSF treatment in homeostatic-enriched (**A**), hybrid (cluster 10) (**B**), DAM-A (**C**), and DAM-B (**D**) MG-sig cluster hubs. Genes upregulated by SCF+G-CSF treatment are shown in red. Genes downregulated by SCF+G-CSF treatment are shown in blue. False discovery rate-corrected *p* values (-log10) are shown on the y-axis. The log2-transformed expression ratios are computed as expression levels in the SCF+G-CSF-treated mice relative to those in the vehicle control group. The log2-transformed expression ratios are plotted on the x-axis. Dotted lines indicate significance thresholds. Note, *p* values are restricted to 300 decimal places. The top 25 differentially-expressed genes are labeled.

**Figure S9. Cluster-specific expression of the topmost differentially-expressed genes upregulated by SCF+G-CSF treatment.** Expression levels of the topmost genes upregulated by SCF+G-CSF treatment are displayed for each MG-sig cluster and Mye-sig cluster in the SCF+G-CSF treatment group (**A**) and in the vehicle control group (**B**). Bubble plots: the diameters of the circles correspond to the percentage of cells that express a given gene while the color intensities of the circles correspond to the magnitude of expression. Note, the differentially expressed genes are more highly expressed in Mye-sig clusters than in MG-sig clusters.

**Figure S10. Microglial cell-selective genes are comparably expressed in S100a8/9-positive and S100a8/9-negative MG-sig cells.** S100a8/9**-**positive cells in MG-sig clusters co-express several microglial gene markers (Cx3cr1, Gpr34, P2ry12, Fcrls, Olfml3, and Tmem119) at comparable levels to S100a8/9-negative cells. Bubble plots: the diameters of the circles correspond to the percentage of cells that express a given gene while the color intensities of the circles correspond to the magnitude of expression.

**Figure S11. S100a8/9-positive MG-sig and Mye-sig cells highly and differentially express the topmost differentially-expressed genes upregulated by SCF+G-CSF treatment.** The topmost 25 genes upregulated by SCF+G-CSF treatment are also co-expressed and increased in S100a8/9**-**positive MG-sig cells (**A**) and S100a8/9-positive Mye-sig cells (**B**). Genes upregulated in S100a8/9-positive cells relative to S100a8/9-negative cells are shown in red. False discovery rate-corrected *p* values (-log10) are shown on the y-axis. The log2-transformed expression ratios are computed as expression levels in S100a8/9-positive cells relative to those in S100a8/9-negative cells. The log2-transformed expression ratios are plotted on the x-axis. Dotted lines indicate significance thresholds.

**Figure S12: Genes identified by network analysis to have direct functional connections to S100a8/9 are largely upregulated in S100a8/9-positive MG-sig cells and S100a8/9-positive Mye-sig cells.** The S100a8/9-linked functional gene set identified by network analysis is largely co-expressed and dramatically upregulated in S100a8/9^+^ MG-sig cells (**A**) and S100a8/9^+^ Mye-sig cells (**B**) compared to S100a8/9-negative MG-sig cells and S100a8/9-negative Mye-sig cells, respectively. Genes upregulated in S100a8/9^+^ MG-sig cells and S100a8/9^+^ Mye-sig cells are shown in red. False discovery rate-corrected *p* values (-log10) are shown on the y-axis. The log2-transformed expression ratios are computed and are plotted on the x-axis. Dotted lines indicate significance thresholds.

**Figure S13. Gene ontology analysis of genes with direct functional connections to S100a8/9.** Gene ontology analysis of genes directly linked with S100a8/9, i.e., direct neighbors as indicated in our network analysis (Fig. 9C), identify associated functions including regulation of the immune system, responses to stress and inflammation, and leukocyte migration and aggregation. These biological functions are largely comparable to analyses of all differentially expressed genes following SCF+G-CSF treatment (Fig. S7).
