## Supplemental Data File Legends for "Single cell RNA sequencing reveals immunomodulatory effects of stem cell factor and granulocyte colony-stimulating factor treatment in the brains of aged APP/PS1 mice"

**Supplemental Data Files**

**Supplemental Data File 1.** Mean expression (log2) of all genes organized by cluster. Rows correspond to individual genes. Columns correspond to individual clusters.

**Supplemental Data File 2.** A complete list of enriched terms from the GO Biological Process and Reactome databases is provided from separate analyses on all genes and on those functionally connected to S100a8/9 as identified by network analysis. Results using the two gene sets are presented in separate tabs.

**Supplemental Data File 3.** A list is provided of the genes differentially expressed in SCF+G-CSF treatment vs. vehicle controls that are common to both MG-sig and Mye-sig clusters. Genes included in the list show ≥ 2-fold change in SCF+G-CSF treatment compared to vehicle controls with an FDR-corrected *p*-value < 0.01. Differential expression values and adjusted *p* values are provided.
